# Excessive Firing of Dyskinesia-Associated Striatal Direct Pathway Neurons is Gated By Dopamine and Excitatory Synaptic Input

**DOI:** 10.1101/2022.10.31.514594

**Authors:** Michael B. Ryan, Allison E. Girasole, Matthew M. McGregor, Rea Brakaj, Ronald F. Paletzki, Charles R. Gerfen, Alexandra B. Nelson

**Affiliations:** Neuroscience Graduate Program, UCSF, San Francisco, CA 94158, USA; Kavli Institute for Fundamental Neuroscience, UCSF, San Francisco, CA 94158, USA; Weill Institute for Neurosciences, UCSF, San Francisco, CA 94158, USA; Department of Neurology, UCSF, San Francisco, CA 94158, USA; Laboratory of Systems Neuroscience, National Institute of Mental Health, Bethesda, Maryland 20892, USA

## Abstract

The striatum integrates dopaminergic and glutamatergic inputs to select preferred versus alternative actions, but the precise mechanisms remain unclear. One way to study action selection is when it breaks down. Here, we explored the cellular and synaptic mechanisms of levodopa-induced dyskinesia (LID), a complication of Parkinson’s disease therapy characterized by involuntary movements. We used an activity-dependent tool (FosTRAP) in conjunction with a mouse model of LID to investigate functionally distinct subsets of striatal direct pathway medium spiny neurons (dMSNs). *In vivo*, levodopa differentially activates dyskinesia-associated (TRAPed) dMSNs compared to other dMSNs. This activation is likely to be driven by two cellular mechanisms we identified through *ex vivo* electrophysiology: higher sensitivity to dopamine and stronger excitatory input from the motor cortex and thalamus. Together, these findings suggest how intrinsic and synaptic properties of heterogeneous dMSN subpopulations integrate to support action selection.

## Introduction

The striatum coordinates many behaviors, ranging from locomotion to reward-based decision making. Within the striatum, two canonical classes of neurons, direct and indirect pathway medium spiny neurons (dMSNs and iMSNs, respectively), are distinguished by their projection targets and dopamine receptor expression (Gerfen et al., 1990). According to classical models of basal ganglia function, dMSNs promote movement, while iMSNs suppress it. Accumulated evidence suggests balanced activity (co-activation) of these two populations supports normal action selection (Barbera et al., 2016; Cruz et al., 2022; Cui et al., 2013; Jin et al., 2014; Mink, 1996). An imbalance may contribute to impaired action selection, resulting in movement or cognitive disorders (DeLong, 1990; McGregor and Nelson, 2019). However, a key question in the field is how heterogeneous subpopulations within these two pathways contribute to action selection in health and disease.

One such disorder is levodopa-induced dyskinesia (LID), a complication of Parkinson’s Disease in which treatment with the dopamine precursor levodopa results in abnormal involuntary movements. Work in nonhuman primate models of Parkinson’s Disease has identified abnormal striatal activity in LID (Liang et al., 2008; Singh et al., 2015), and cell type-specific approaches in rodent models indicate LID is accompanied by increased dMSN and decreased iMSN activity, consistent with classical models (Parker et al., 2018; Ryan et al., 2018; Sagot et al., 2018). In keeping with the idea that direct and indirect pathways may contain functional subdivisions, we and others have observed considerable variability in how MSNs respond to levodopa and how their firing relates to dyskinesia *in vivo* (Fieblinger et al., 2018; Liang et al., 2008; Parker et al., 2018; Ryan et al., 2018). We identified a subpopulation of dMSNs *in vivo* with exceptionally high firing rates in response to levodopa, which in turn correlate with dyskinesia severity (Ryan et al., 2018). In a parallel study, we used an activity-dependent transgenic mouse line (FosTRAP) (Guenthner et al., 2013) to capture highly active neurons during a single episode of LID; we found that re-activation of this subpopulation could drive dyskinesia in the absence of levodopa (Girasole et al., 2018).

Here, we used FosTRAP to explore the mechanisms that drive LID. We found that a subpopulation of dMSNs labeled by FosTRAP show exceptionally high levodopa-evoked firing, which correlates with dyskinesia severity on a moment-to-moment basis. To identify the synaptic changes that might underlie this aberrant firing pattern, we used Cre-dependent rabies tracing, optogenetics, and *ex vivo* physiology, finding that this subpopulation of dMSNs has stronger excitatory synaptic input than neighboring dMSNs or iMSNs, and is more sensitive to dopamine signaling. Together, these findings identify specific cellular mechanisms that underlie heterogeneous responses to levodopa, driving therapeutic effects as well as dyskinesias.

## Results

To investigate the physiological drivers of LID, we captured LID-associated striatal neurons using FosTRAP in a toxin-based mouse model of PD and LID (Cenci and Lundblad, 2007) (Figure 1A). The left medial forebrain bundle of FosTRAP;Ai14 (TRAP) mice was injected with the dopaminergic neurotoxin 6-hydroxydopamine (6-OHDA), which caused loss of midbrain dopamine neurons (Figure S1A), ipsilateral rotational bias and reduced movement velocity (Figures S1B, D). Several weeks later, mice began daily levodopa treatment, resulting in contralateral rotational bias, increased movement velocity, and robust LID (Figures 1B-C; S1C-D). LID-associated neurons were captured (“TRAPed”) one week into the treatment course with coadministration of levodopa and 4-OH tamoxifen, driving Cre-dependent expression of tdTomato (Figure 1D) (Girasole et al., 2018).

**Figure 1.**
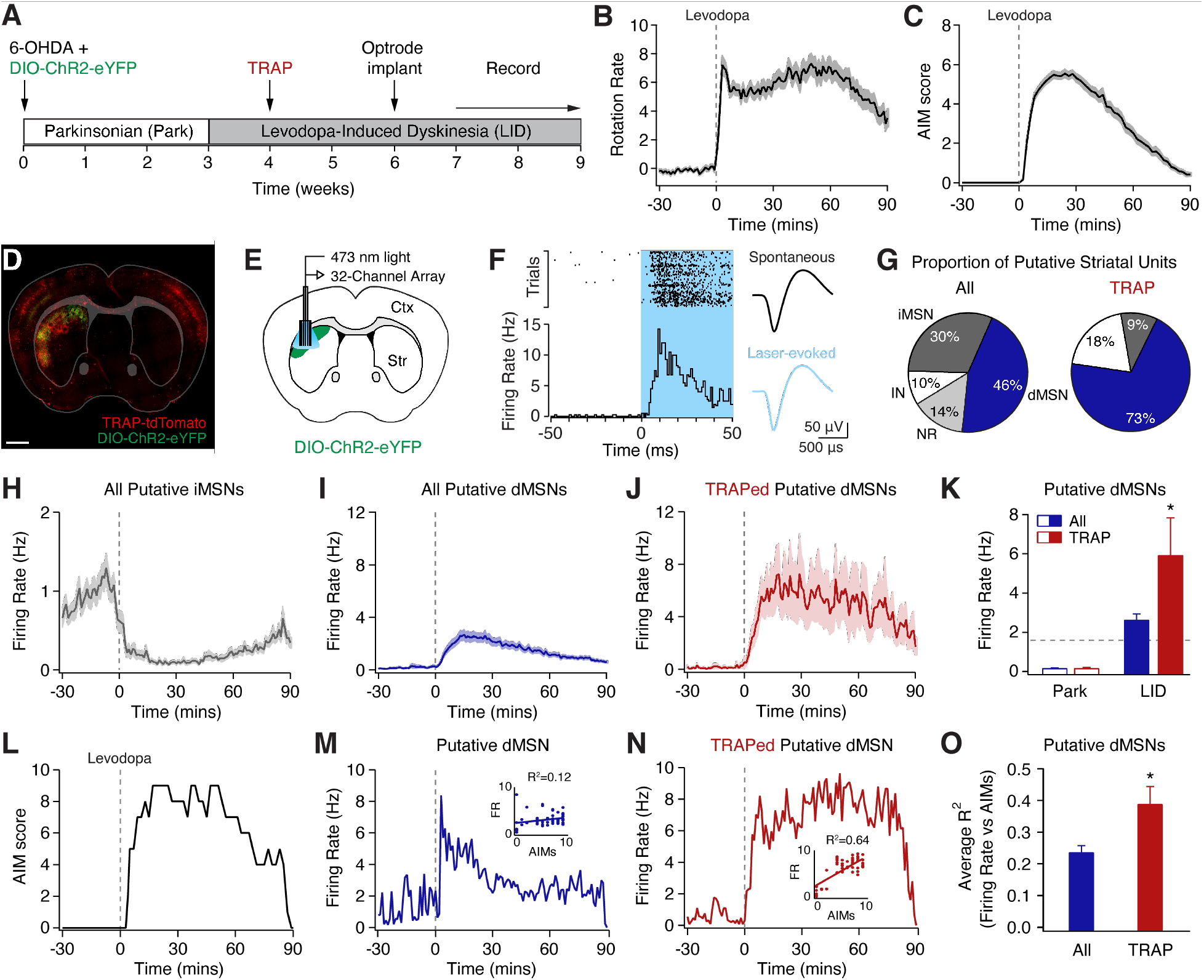
Optogenetically Identified TRAPed Striatal Neurons Show Differential Firing Responses to Levodopa *In Vivo*. TRAPed striatal single-units were recorded in freely moving parkinsonian mice using an optogenetic labeling approach. **(A)** Experimental timeline. **(B-C)** Rotation rate (B) (contralesional-ipsilesional rotations per minute and dyskinesia (C) (quantified with Abnormal Involuntary Movement, AIM score) of parkinsonian mice, aligned to levodopa injection at t=0 (N=14). **(D-E)** DIO-ChR2-eYFP and optrode array in the dorsolateral striatum of a FosTRAP;Ai14 mouse. Representative postmortem histology (D) and coronal schematic (E). **(F)** Representative optogenetically labeled TRAPed striatal unit. Left: peri-event raster (top) and histogram (bottom) showing spiking in response to laser (blue box). Right: average spontaneous (top) and laser-evoked (bottom) waveforms. **(G)** Proportion of all (left; n=268, N=14) and optically labeled TRAPed (right; n=11, N=8) striatal units, including putative interneurons (IN), direct pathway (dMSN), indirect pathway (iMSN), and no response (NR) striatal units. **(H-J)** Average firing rate of putative iMSNs (n=81, N=14), dMSNs (n=117, N=14), and optically labeled TRAPed dMSNs (n=8, N=6), aligned to levodopa injection at t=0. **(K)** Average firing rate of all putative dMSNs and optogenetically labeled TRAPed dMSNs in parkinsonian mice before (Park) and after levodopa administration (LID). Dotted line represents the firing rate of optogenetically labeled dMSNs from healthy controls (from Ryan et al., 2018). * p<0.05, Wilcoxon Rank Sum Test **(L-N)** Data from a single recording session, including **(L)** Dyskinesia score, **(M)** firing rate of a dMSN, and **(N)** firing rate of an optogenetically labeled TRAPed dMSN. Insets: firing rate vs dyskinesia score. **(O)** Average correlation (R^2^) of firing rate to dyskinesia for all dMSNs vs and TRAPed dMSNs. n=single units, N=mice. All data presented as mean ± SEM. * p<0.05, Wilcoxon Rank Sum Test. See also Figure S1.

### TRAPed Direct Pathway Neurons Respond Differentially to Levodopa In Vivo

Levodopa changes both firing rate and pattern in MSNs in animal models of PD (Liang et al., 2008; Ryan et al., 2018; Singh et al., 2015). To compare the firing of TRAPed cells to other striatal neurons, we performed optogenetically labeled single-unit recordings in the dorsolateral striatum (DLS) of freely moving parkinsonian mice (Figure 1E). TRAPed neurons were optically identified by their short-latency light-evoked responses (Figure 1F)(Kravitz et al., 2013). In line with previous findings, we observed a variety of responses to levodopa in striatal neurons, including decreased firing (Figures S1E-F), increased firing (Figures S1G-H), and no significant change (Figure S1I). Based on previous optically labeled recordings in a similar model, we classified these units as putative iMSNs, dMSNs, or NR (no response) units (Figures 1G-I)(Ryan et al., 2018). We also identified putative interneurons based on spike waveform (Figures S1K-L) (Barnes et al., 2005; Gage et al., 2010). We found TRAPed neurons were highly enriched for dMSNs as compared to the overall population (Figure 1G), as predicted by previous histological analysis (Girasole et al., 2018). In the parkinsonian state, TRAPed neurons and other dMSNs had comparably low firing rates, but in response to levodopa, TRAPed neurons achieved very high firing rates (Figures 1I-K). These results indicate TRAPed neurons represent a distinct subpopulation of striatal MSNs.

We next examined the firing dynamics of TRAPed dMSNs. Consistent with previous work, we found the firing of some, but not all, dMSNs correlates with dyskinesia severity (Figures 1L-N)(Ryan et al., 2018). Given the causal link between activation of TRAPed dMSNs and dyskinesia (Girasole et al., 2018), we hypothesized their firing would be more tightly correlated to dyskinesia severity. Indeed, we found that as compared to all dMSNs, the firing of TRAPed dMSNs was more highly correlated with dyskinesia (Figures 1O; S1J). Together, these findings indicate that the firing of TRAPed dMSNs is not only more sensitive to levodopa, but correlates with dyskinesia on a moment to moment basis.

### TRAPed Direct Pathway Neurons Are More Sensitive to Dopamine Signaling

Our *in vivo* findings suggested TRAPed dMSNs are a subpopulation with distinct physiological responses to levodopa. These responses might be mediated by differences in intrinsic or synaptic properties, or sensitivity to dopamine signaling. To address these possibilities, we made *ex vivo* whole-cell patch-clamp recordings from identified DLS TRAPed dMSNs, unTRAPed dMSNs, and iMSNs, using FosTRAP;Ai14;D2-GFP mice (Figures 2A-B). In current-clamp recordings, we found that basic properties did not differ between the three cell types (Figures 2C-F; Supplemental Table 1). We hypothesized that D1R activation would increase the excitability of TRAPed dMSNs based on previous studies showing similar effects in healthy animals (Hernandez-Lopez et al., 2000; Lahiri and Bevan, 2020; Planert et al., 2013). The D1R agonist SKF81297 increased the excitability of TRAPed dMSNs, while unTRAPed iMSNs and dMSNs showed no significant change (Figures 2G-I; Supplemental Table 2). Thus, differential sensitivity to dopamine by TRAPed dMSNs is one potential mechanism for their high levodopa-evoked firing rates *in vivo*.

**Figure 2:**
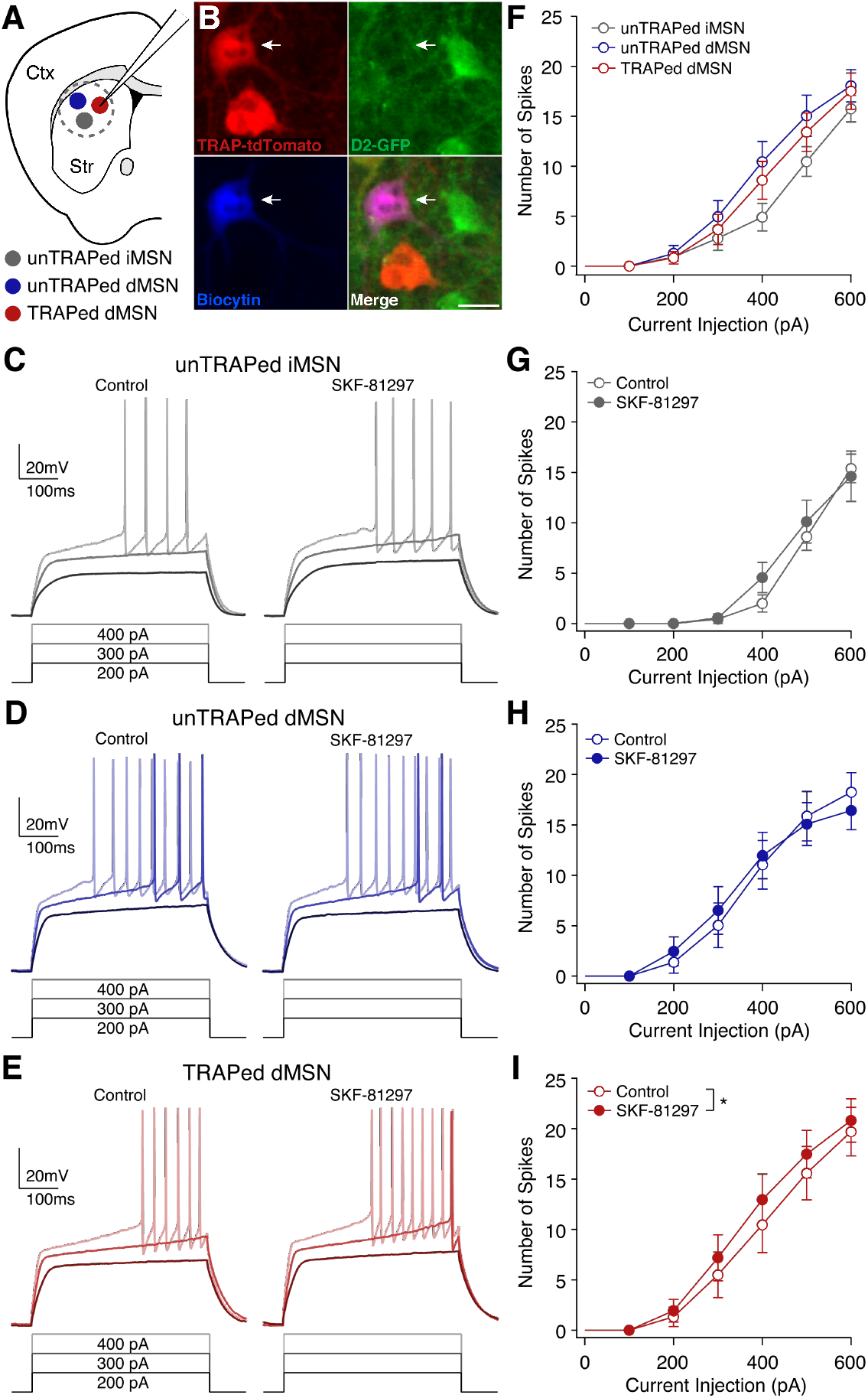
Activation of Dopamine D1 Receptors Enhances the Excitability of TRAPed, but not unTRAPed, dMSNs. Striatal neurons were targeted for *ex vivo* whole-cell recordings in coronal brain slices from parkinsonian and levodopa-treated FosTRAP;Ai14;D2-GFP (FAD) mice. **(A)** Cartoon showing recordings of unTRAPed iMSNs (gray), unTRAPed dMSNs (blue), and TRAPed dMSNs (red) in the dorsolateral striatum. **(B)** Histological image showing a biocytin-filled TRAPed dMSN targeted for recording (arrow). Tissue shows expression of FosTRAP;Ai14 (tdTomato, red), D2R (GFP, green) and biocytin (blue). Scale bar = 20 µm. **(C-E)** Representative voltage responses to current injections before (left) and 10-15 minutes after bath application of the D1R-agonist, SKF-81297 (right) for a unTRAPed iMSN (C), unTRAPed dMSN (D), and TRAPed dMSN (E).**(F)** Average current-response curves for unTRAPed iMSNs (gray, n=17, N=11), unTRAPed dMSNs (blue, n=17, N=13), and TRAPed dMSNs (red, n= 22, N=14). **(G-I)** Current-response curves before (Control) and 10-15 minutes after SKF-81297 for unTRAPed iMSNs (n=9, N=6; G), unTRAPed dMSNs (n=11, N=9; H), and TRAPed dMSNs (n=14, N=10; I). n=cells, N=mice. Data presented as mean ± SEM. See also Tables 1-2.

### Monosynaptic Rabies Tracing Reveals Reduced Number of Excitatory Inputs Onto TRAPed Direct Pathway Neurons in LID

Striatal MSNs are highly dependent on excitatory synaptic input to drive spiking (Wickens and Wilson, 1998). To determine whether the number or distribution of synaptic inputs to TRAPed dMSNs might contribute to their *in vivo* firing dynamics and more broadly to LID, we performed Cre-dependent rabies tracing (Wickersham et al., 2007) from the DLS (Figure 3A). We used D1-Cre, A2a-Cre, and FosTRAP^CreER^ mice to enable comparisons between inputs to dMSNs, iMSNs, and TRAPed striatal neurons, respectively (Figures 3B-C). Striatal “starter” cells and presynaptic rabies-labeled cell bodies were detected, mapped onto the Allen Brain Atlas and quantified by brain region (Figures 3D-G; S2A-D) (Eastwood et al., 2019). To quantify the relative number of presynaptic neurons, we first calculated the relative number of presynaptic neurons to co-infected striatal starter cells (referred to as “number”, Figure S2D) across healthy, parkinsonian, and LID conditions (Figures 3H-K). To quantify shifts in the distribution of presynaptic neurons across the brain, we also calculated the relative proportion of presynaptic neurons in one area versus the total number of labeled extrastriatal presynaptic neurons (referred to as “proportion” (Figures S2E-H). In the healthy condition, monosynaptic inputs onto dMSNs and iMSNs showed a similar number of presynaptic neurons (Figure 3H), the majority of which derive from the ipsilateral cortex, thalamus, and external segment of the globus pallidus (GPe) (Figures S2E-H), as has been previously reported (Guo et al., 2015; Wall et al., 2013).

**Figure 3.**
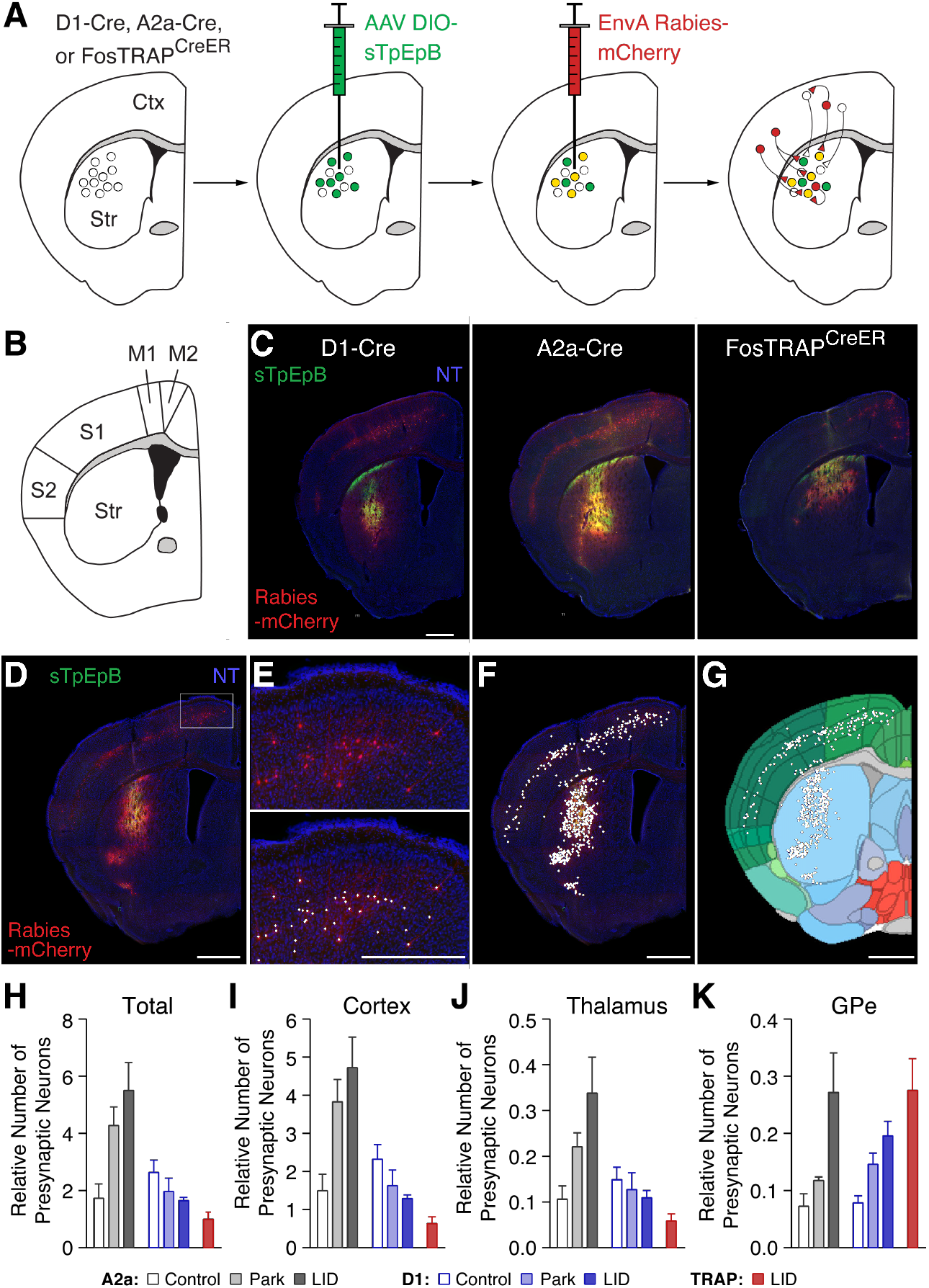
Monosynaptic Rabies Tracing of Striatal Inputs in Healthy, Parkinsonian, and Levodopa-Treated Mice. A dual viral, Cre-dependent strategy was used to label monosynaptic inputs onto direct pathway, indirect pathway, and TRAPed striatal neurons. **(A)** Cartoon of the experimental approach in D1-Cre, A2a-Cre and FosTRAP^CreER^ mice. **(B-C)** Coronal schematic (B) and low magnification histological sections (C) showing helper virus expressing neurons (sTpEpB, green) and rabies-labeled neurons (Rabies-mCherry, red) in D1-Cre (left), A2a-Cre (middle), and FosTRAP^CreER^ (right) mice. DAPI = nuclear stain for visualization. **(D-G)** Quantification of labeled presynaptic cell bodies. (D) Low magnification image of coronal section showing helper (green) and rabies (red) viral injection sites. (E) High magnification of the box in (D), with overlaid points denoting rabies-positive presynaptic cell bodies (bottom). (F-G) Low magnification image showing coronal section (F) and Allan Brain Atlas (G) with overlaid cell detection. **(H-K)** relative number of presynaptic neurons, calculated as the number of extra-striatal rabies labeled neurons divided by the number of co-infected (rabies/helper virus labeled) striatal neurons for all extra-striatal brain regions (H), cortex (I), thalamus (J), and external globus pallidus (K). A2a: Control, N=6, Park, N=4, LID, N=4; D1: Control, N=9, Park, N=10, LID, N=6; TRAP: LID, N=6. Data presented as mean ± SEM. N=animals. Scale bars = 1 mm. See also Figure S2.

In parkinsonian mice, we observed opposing changes in synaptic inputs onto iMSNs and dMSNs. Compared to healthy mice, iMSNs in parkinsonian mice showed an increase in the number of cortical presynaptic neurons, with no significant change in the number of thalamic or GPe input neurons (Figures 3I-K), leading to an overall *increase* in the proportion of cortical inputs to iMSNs (Figure S2E). In contrast, dMSNs showed no change in the number of cortical and thalamic presynaptic neurons, but an increase in the number of presynaptic GPe neurons (Figure 3I-K). This increase in GPe inputs led to a *decrease* in the overall proportion of cortical inputs to dMSNs in parkinsonian mice (Figure S2E). Levodopa-treated parkinsonian animals showed similar overall patterns of monosynaptic inputs to those seen in untreated parkinsonian mice. In iMSNs, there was no significant change in the number of cortical or thalamic presynaptic neurons (Figures 3I-J). However, there was an increase in the number of presynaptic GPe neurons compared to untreated parkinsonian mice (Figure 3K). In dMSNs, we observed no significant changes in the pattern of monosynaptic inputs in levodopa-treated, compared to untreated, parkinsonian animals (Figures 3I-K). Together, these results suggest that chronic levodopa administration does not markedly change the distribution of monosynaptic inputs onto canonical MSN subclasses in parkinsonian mice.

To investigate the possible synaptic drivers of aberrant activity of TRAPed neurons during LID, we next compared the distribution of monosynaptic inputs onto TRAPed neurons versus all dMSNs, focusing on levodopa-treated parkinsonian mice. We hypothesized that TRAPed neurons might receive inputs from a larger number of excitatory cortical and thalamic neurons. We found that the total number of presynaptic neurons brain-wide was similar onto TRAPed neurons and all dMSNs (Figure 3H). However, contrary to our hypothesis, TRAPed neurons had a smaller number of inputs from excitatory sources (thalamus and cortex) than did dMSNs more broadly (Figures 3I, J). This reduction in the number of presynaptic cortical neurons was observed across most motor and somatosensory cortices, leading to a striking reduction in the proportion of cortical inputs (Figures S2E, H). Taken together, these findings suggest that chronic changes in dopamine produce opposing changes in synaptic inputs onto iMSNs and dMSNs, with a marked loss in the number of corticostriatal neurons synapsing onto TRAPed neurons (and dMSNs) in LID. In isolation, this observation is unlikely to explain the high levodopa-evoked firing of TRAPed neurons *in vivo*.

### Synaptic Physiology Shows Enhanced Motor Cortical and Thalamic Excitatory Input Onto TRAPed Direct Pathway Neurons in LID

Though rabies tracing showed structural differences in excitatory synaptic input onto TRAPed dMSNs, this approach may fail to capture synaptic function. To directly compare the strength of excitatory synapses onto neighboring TRAPed dMSNs, unTRAPed dMSNs, and unTRAPed iMSNs, we again used *ex vivo* slice recordings. Based on the tight correlation of TRAPed dMSN spiking to dyskinesia *in vivo*, we hypothesized that excitatory input would be increased onto TRAPed neurons. We tested this hypothesis with several electrophysiological measures that reflect pre and postsynaptic function. To assess presynaptic changes, we first measured miniature excitatory postsynaptic currents (mEPSCs), which were increased in frequency, but not amplitude, in TRAPed dMSNs (Figures 4A-B; S3A-B), consistent with a possible increase in presynaptic number or probability of release. To assess for postsynaptic changes, we next measured electrically evoked EPSCs. We found no differences in the AMPA:NMDA ratio (Figures S3C-D). However, in the same recordings we found a marked decrease in the paired pulse ratio (PPR) in TRAPed dMSNs as compared to unTRAPed dMSNs or iMSNs (Figures 4C-D). Together, these results suggest a higher probability of release at excitatory synapses onto TRAPed dMSNs compared to other MSNs.

**Figure 4:**
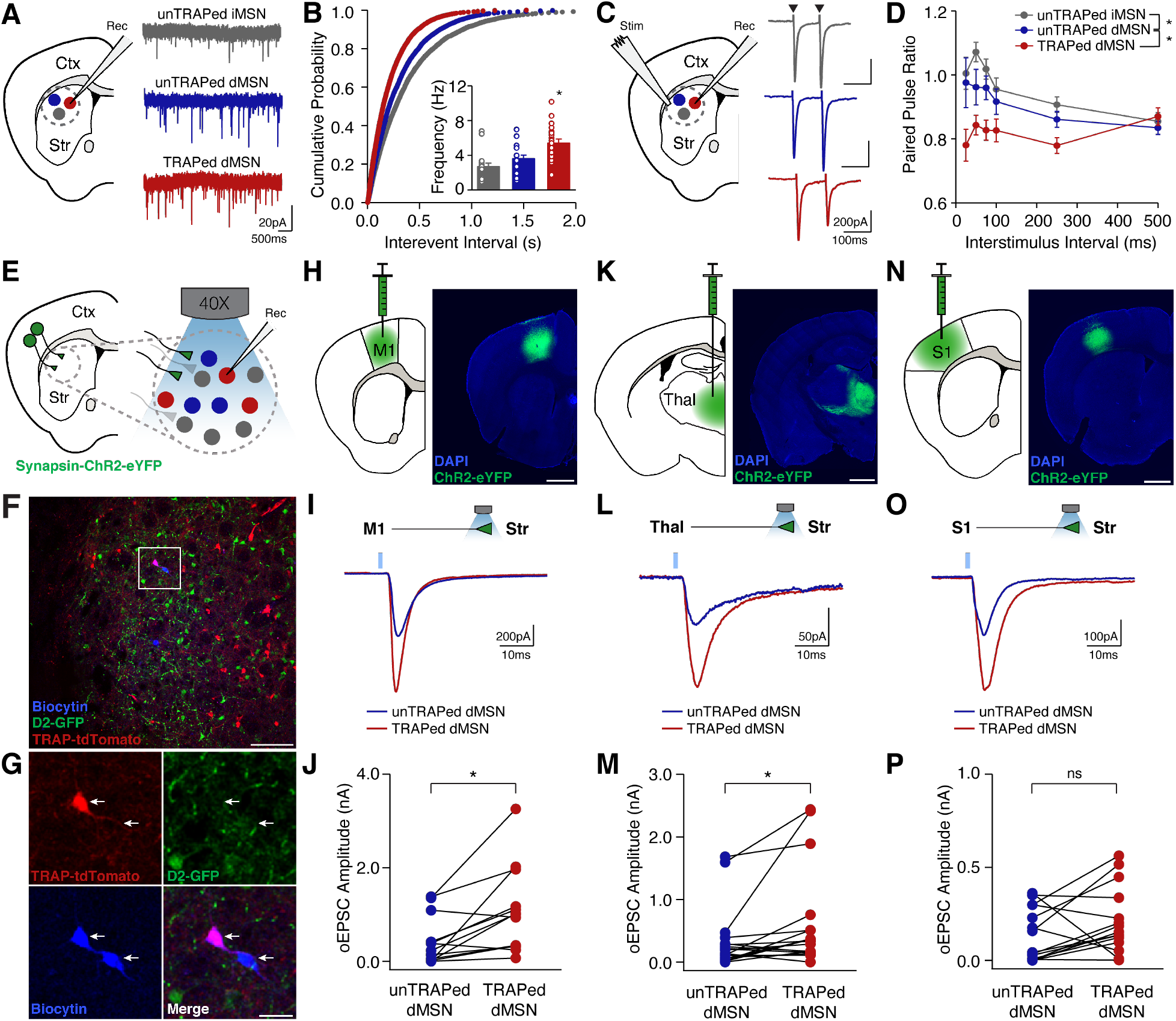
Increased Presynaptic Excitatory Transmission onto TRAPed dMSNs. Excitatory inputs to striatal dMSNs were compared in ex vivo brain slices from the dorsolateral striatum of FAD mice **(A-D). (A)** Coronal schematic (left) and representative current traces (right) from unTRAPed iMSNs, unTRAPed dMSNs, and TRAPed dMSNs in the presence of picrotoxin and tetrodotoxin to isolate miniature excitatory postsynaptic currents (mEPSCs). **(B)** Cumulative probability distribution of mEPSC frequencies. Inset: average mEPSC frequencies (unTRAPed iMSNs: n=21, N=8; unTRAPed dMSNs: n=20, N=7; TRAPed dMSNs: n=20, N=8). **(C)** Coronal schematic (left) and representative traces (right) from unTRAPed iMSNs, unTRAPed dMSNs, and TRAPed dMSNs in response to local electrical stimulation (arrowheads) in the presence of picrotoxin, to isolate evoked EPSCs. (D) Summary of the paired pulse ratio in unTRAPed iMSNs (n=17, N=8), unTRAPed dMSNs (n=18, N=9), and TRAPed dMSNs (n=22, N=9). **(E-P)** The strength of major excitatory inputs was compared across unTRAPed and TRAPed dMSNs using an optogenetic approach, n=pairs, N=mice. (**E)** Cartoon of recording configuration. **(F)** Low magnification image of dorsolateral striatum, showing TRAPed neurons (red), D2R-expressing neurons (green), and a pair of biocytin-filled neurons (blue). Scale bar = 100 µm. **(G)** High magnification of section in (F), showing a pair of neighboring, sequentially recorded TRAPed and unTRAPed dMSNs (arrows). Scale bar = 25 µm. **(H-P)** Optical activation of inputs from primary motor cortex (H-J, M1, n=19, N=4), thalamus (K-M, (Thal, n=15, N=5), and primary somatosensory cortex (N-P, S1, n=13, N=7). **(H, K, N)** Coronal schematics (left) and postmortem histology (right) showing viral expression of ChR2-eYFP. Scale bar = 1 mm. **(I, L, O)** Representative examples of optically-evoked EPSCs (oEPSCs) for an unTRAPed and TRAPed dMSN (bottom). **(J, M, P)** Average oEPSC amplitude at 4mW for unTRAPed and TRAPed dMSNs. * p<0.05, Wilcoxon Rank Sum Test. n=cells, N=mice. Data presented as mean ± SEM.

While these findings suggest greater strength of excitatory input onto TRAPed dMSNs, they do not identify the specific source. We next assessed the strength of several key inputs to the DLS using an optical approach (Figures 4E-G). We injected hSyn-ChR2 into primary motor cortex (M1), somatosensory cortex (S1) and thalamus (Figures 4H, K, N). Comparing paired recordings, we found optically evoked EPSCs from M1 and thalamus were larger on TRAPed than unTRAPed dMSNs (Figures 4H-M; S3E-H). Optically evoked S1 EPSCs were not statistically different between groups (Figures 4N-P; S3I-J). These results indicate TRAPed dMSNs receive differential excitatory synaptic input. Taken together, a selective enhancement in excitatory inputs combined with a dopamine-dependent increase in excitability might underlie the excessive levodopa-evoked firing of TRAPed dMSNs *in vivo*.

## Discussion

Here we have used FosTRAP in a mouse model of levodopa-induced dyskinesia (LID) to explore the mechanisms and consequences of heterogeneity within one of the canonical striatal pathways: the direct pathway. While previous work has described striatal subpopulations, based on molecular identity, anatomical location, inputs or outputs (Cazorla et al., 2014; Flaherty and Graybiel, 1991; Gokce et al., 2016; Hintiryan et al., 2016; Martin et al., 2019; Saunders et al., 2018; Stanley et al., 2020), others have shown relative homogeneity within pathways (Peng et al., 2021), and connecting these features to functional roles *in vivo* has been more challenging. Using FosTRAP to label neurons whose activation is known to cause dyskinesia, we were able to test whether *in vivo* or *ex vivo* changes associated with LID are differentially expressed between dMSNs.

We found that *in vivo*, optically identified TRAPed neurons were enriched for a subpopulation of DLS dMSNs previously identified by their firing and functional features (Ryan et al., 2018). Their transition from low to exceptionally high levodopa-evoked firing rates may explain the large levodopa-activated dMSN ensembles seen in a recent calcium imaging study in parkinsonian mice (Maltese et al., 2021). While TRAPed dMSNs represent a subset of all dMSNs, our new findings suggest they both correlate with and cause LID in the mouse model. What are the underlying mechanisms that produce this distinct phenotype? There are two major possibilities: intrinsic excitability and/or synaptic input.

The *in vivo* properties of TRAPed dMSNs could be explained by increased intrinsic excitability, making them more likely to spike to a given synaptic input. Previous work has found increases in dMSN excitability in chronically parkinsonian mice, and only partial restoration in levodopa-treated animals (Fieblinger et al., 2014). In fact, a recent study found heterogeneity in intrinsic excitability between MSNs in the mouse model of LID (Fieblinger et al., 2018). However, we found that basal excitability was quite similar between TRAPed and unTRAPed dMSNs. This matches well with our observation that *in vivo*, dMSNs show uniformly low firing rates in the parkinsonian condition – only upon levodopa administration are the differences between dMSNs apparent. Indeed, we found that as compared to unTRAPed dMSNs, TRAPed dMSNs showed a modest increase in excitability in response to dopamine signaling. This differential response might be mediated by increased D1 receptors or amplified downstream signaling regulating excitability in TRAPed dMSNs (Gerfen et al., 2002; Heiman et al., 2014; Jenner, 2008; Lahiri and Bevan, 2020). As levodopa treatment would be expected to increase local dopamine signaling, this might amplify the responses of TRAPed dMSNs to their synaptic inputs.

In addition, the excitatory synaptic inputs to TRAPed dMSNs may be enhanced, particularly from sensorimotor areas that are likely to drive voluntary (and involuntary) movements. Previous studies have used slice electrophysiology to examine excitatory synaptic inputs onto striatal neurons in rodent models of PD and LID, finding on the one hand an increase in synaptic strength onto MSNs overall (Bagetta et al., 2012), and on the other a decrease in synaptic input onto dMSNs specifically (Fieblinger et al., 2014). Here we used an anatomical assay (monosynaptic rabies tracing) and a physiological assay (slice electrophysiology) to investigate alterations in synaptic input onto TRAPed MSNs. While rabies tracing revealed rearrangements of presynaptic inputs onto dMSNs and iMSNs between the healthy and parkinsonian (with and without levodopa) conditions, this approach did not find increases in the relative number of excitatory presynaptic neurons targeting dMSNs or TRAPed dMSNs. In fact, the overall reduction in presynaptically labeled neurons may relate to the reductions in spine density seen on dMSNs in the mouse model of LID (Fieblinger et al., 2014). However, slice electrophysiology indicated these inputs were much stronger in TRAPed dMSNs versus their unTRAPed dMSN neighbors. Why are these results seemingly discordant? With rabies tracing, we quantified the number of presynaptic neurons projecting to a cell type of interest (dMSNs, iMSNs, or TRAPed neurons). However, a single presynaptic neuron may make profuse contacts onto a single or multiple postsynaptic targets (Cho et al., 2002), explaining the markers of high synaptic connectivity (reduced paired pulse ratio, high mEPSC frequency) we observed in TRAPed neurons, as well as the increased amplitude of optically-evoked responses onto TRAPed neurons. This pattern of connectivity could drive selective but exceptionally high firing during dyskinesia we observed in TRAPed dMSNs *in vivo*. Using similar methods, another group has identified increased corticostriatal connectivity in the context of cocaine sensitization (Wall et al., 2019).

There are several limitations to our study. As FosTRAP relies on cFos activation, which is minimal in the healthy and parkinsonian striatum, we were not able to compare the same subpopulation of neurons across healthy, parkinsonian but levodopa-naïve, and chronically levodopa-treated states. Instead, we compared the properties of TRAPed neurons before and after acute dopaminergic signaling, and to neighboring unTRAPed neurons in the same state. TRAPed MSNs are predominantly dMSNs, but iMSNs contribute to dyskinesia, as well (Alcacer et al., 2017). We focused on excitatory input, as this is a key driver of MSN firing, but alterations in the strength or sources of inhibition (Glajch et al., 2016; Taverna et al., 2008; Tepper et al., 2018), as well as cholinergic signaling (Bordia and Perez, 2019; Shen et al., 2016; Won et al., 2014), might also potently shape MSN firing during LID. In fact, rabies tracing suggested that inhibitory GPe-MSN inputs were increased in parkinsonian/levodopa-treated mice, and this was even more pronounced in TRAPed neurons. Future work will need to address the role of these other inputs in shaping striatal activity during LID.

Finally, the differences we observed between TRAPed dMSNs and their neighbors may be related to heterogeneity present in the healthy striatum (fixed factors), or may be related to dopamine-dependent changes that are unevenly distributed across the striatum (plasticity). In fact, healthy striatal subpopulations, defined by receptor expression, specific cortical inputs, or local connectivity, may be differentially vulnerable to forms of plasticity known to occur in response to chronic dopamine depletion and dopamine replacement. Together, these alterations in excitability or the strength and pattern of inputs may subserve homeostasis in the healthy brain, or aid compensation in disease states, but when pushed to their limits, drive aberrant circuit function and behavior as are seen in LID.

