## Supplemental Figures and Text for "Excessive Firing of Dyskinesia-Associated Striatal Direct Pathway Neurons is Gated By Dopamine and Excitatory Synaptic Input"

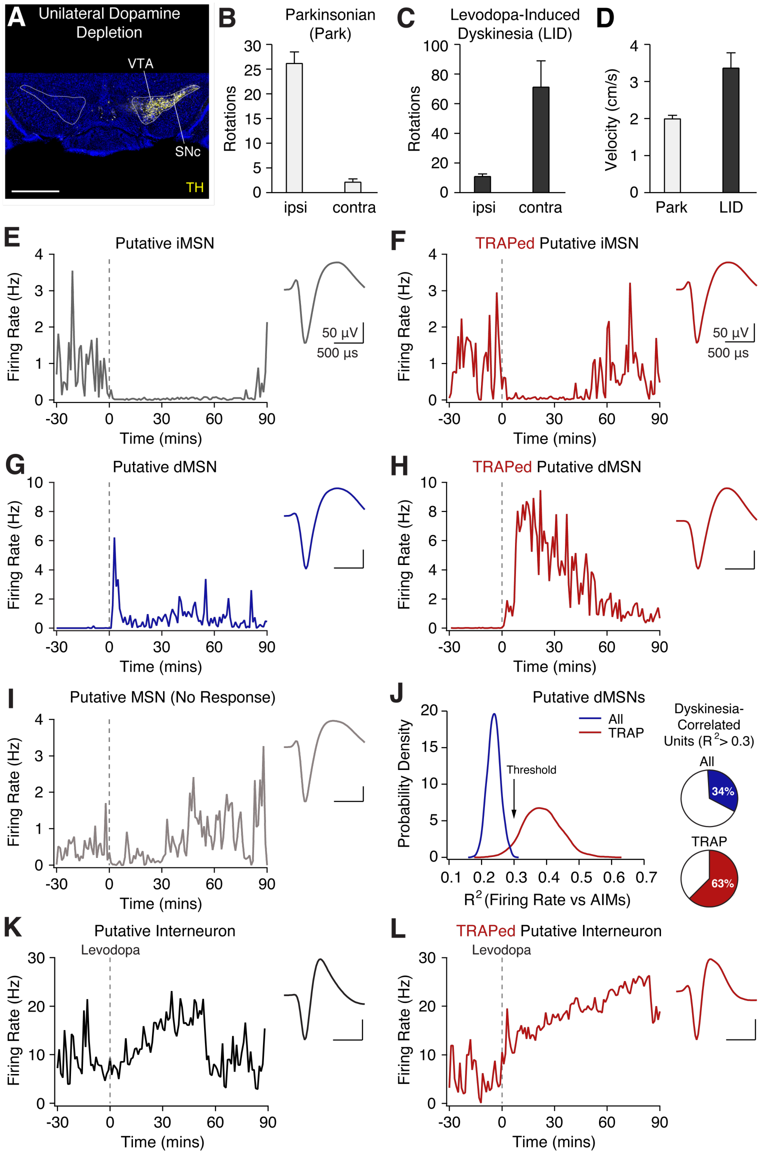


**Figure S1: Optogenetically Identified TRAPed Striatal Neurons Show Differential Responses to Levodopa In Vivo. Related to Figure 1.**

TRAPed striatal single-units were recorded in freely moving parkinsonian mice using an optogenetic labeling approach. **(A)** Representative histological section showing tyrosine-hyrdroxylase (TH) staining in the midbrain of a hemiparkinsonian mouse. Scale bar = 1 mm. (**B-C)** Ipsilesional (ipsi) and contralesional (contra) rotations over 10 minutes in parkinsonian mice before (B) and 20 minutes after levodopa (C). **(D)** Movement velocity of parkinsonian mice before (Park) and 20 minutes after levodopa injection (LID). **(E-I, K-L)** Representative single units from parkinsonian mice treated with levodopa. Left: single unit firing rates, aligned to levodopa administration at t=0. Right: average waveform. **(E,F)** Unlabeled (E) and optogenetically labeled TRAPed (F) putative iMSNs. **(G,H)** Unlabeled (G) and optogenetically labeled TRAPed (H) putative dMSNs. **(I)** Unlabeled MSN with no response to levodopa **(J)** Left: Probability density plot of average correlation ($R^2$) obtained from bootstrapping all putative dMSN (blue) and optogenetically labeled TRAPed putative dMSN data (red). Right: Proportion of all (n=117, N=14) and TRAPed (n=8, N=6) putative dMSNs with a significant correlation ($R^2 > 0.3$) between firing rate and dyskinesia score. **(K,L)** Unlabeled (K) and optogenetically labeled TRAPed (L) putative interneurons.n=single units, N=mice.

| | unTRAPed iMSNs | unTRAPed dMSNs | TRAPed dMSNs | p-value |
|---|---|---|---|---|
| Input Resistance (MOhm) | 79.1 ± 9.5 | 81.1 ± 5.8 | 82.3 ± 7.1 | 0.503 |
| Resting Membrane Potential (mV) | -88.3 ± 1.0 | -89.2 ± 0.9 | -89.4 ± 0.8 | 0.542 |
| Action Potential Threshold (mV) | -39.8 ± 2.7 | -38.9 ± 1.5 | -38.9 ± 1.8 | 0.834 |
| AHP Amplitude (mV) | 15.6 ± 1.1 | 13.8 ± 1.1 | 14.7 ± 1.0 | 0.238 |
| Spike Width (ms) | 1.8 ± 0.1 | 2.0 ± 0.1 | 2.1 ± 0.1 | 0.159 |
| Rheobase (pA) | 406.1 ± 39.6 | 341.2 ± 25.1 | 384.2 ± 7.1 | 0.447 |

**Table 1. Baseline Passive and Active Membrane Properties of MSNs. Related to Figure 2.**
Passive (input resistance and resting membrane potential) and active properties (action potential threshold, after-hyperpolarization amplitude, spike width, and rheobase) of excitability in the absence of dopamine receptor stimulation. Data presented as mean ± SEM.

| | unTRAPed iMSNs | | | unTRAPed dMSNs | | | TRAPed dMSNs | | |
|---|---|---|---|---|---|---|---|---|---|
| | **Control** | **SKF-81297** | **p-value** | **Control** | **SKF-81297** | **p-value** | **Control** | **SKF-81297** | **p-value** |
| Input Resistance (MOhm) | 79.8 ± 13.8 | 91.8 ± 14.5 | 0.383 | 82.0 ± 7.8 | 99.9 ± 13.4 | 0.032 | 89.4 ± 8.7 | 90.4 ± 8.3 | 0.925 |
| Resting Membrane Potential (mV) | -89.8 ± 0.8 | -86.8 ± 3.1 | 0.469 | -90.5 ± 1.1 | -89.0 ± 1.3 | 0.320 | -90.3 ± 1.0 | -90.0 ± 1.6 | 0.791 |
| Action Potential Threshold (mV) | -39.5 ± 3.5 | -38.5 ± 3.0 | 0.313 | -37.5 ± 1.9 | -35.9 ± 1.6 | 0.910 | -38.9 ± 2.3 | -38.7 ± 2.8 | 0.910 |
| AHP Amplitude (mV) | 21.4 ± 3.2 | 20.8 ± 3.3 | 0.148 | 28.8 ± 3.3 | 22.8 ± 2.5 | 0.700 | 27.1 ± 2.3 | 22.9 ± 3.9 | 0.204 |
| Spike Width (ms) | 1.8 ± 0.2 | 2.2 ± 0.7 | 0.313 | 2.0 ± 0.1 | 2.1 ± 0.1 | 0.049 | 2.1 ± 0.6 | 2.2 ± 0.7 | 0.233 |
| Rheobase (pA) | 440.0 ± 51.5 | 405 ± 51.9 | 0.250 | 345.8 ± 29.2 | 312.5 ± 28.3 | 0.125 | **371.4 ± 29.0** | **321.4 ± 31.8** | **0.0078** |

**Table 2. Passive and Active Membrane Properties of MSNs in Response to the D1-receptor specific agonist, SKF-81297. Related to Figure 2.**
Passive (input resistance and resting membrane potential) and active properties (action potential threshold, after-hyperpolarization amplitude, spike width, and rheobase) of excitability before (Control) and 10-15 minutes after bath application of a D1-specific agonist (SKF-81297). Bolded values represent significant p-values, following Bonferroni correction for multiple comparisons. Data presented as mean ± SEM.

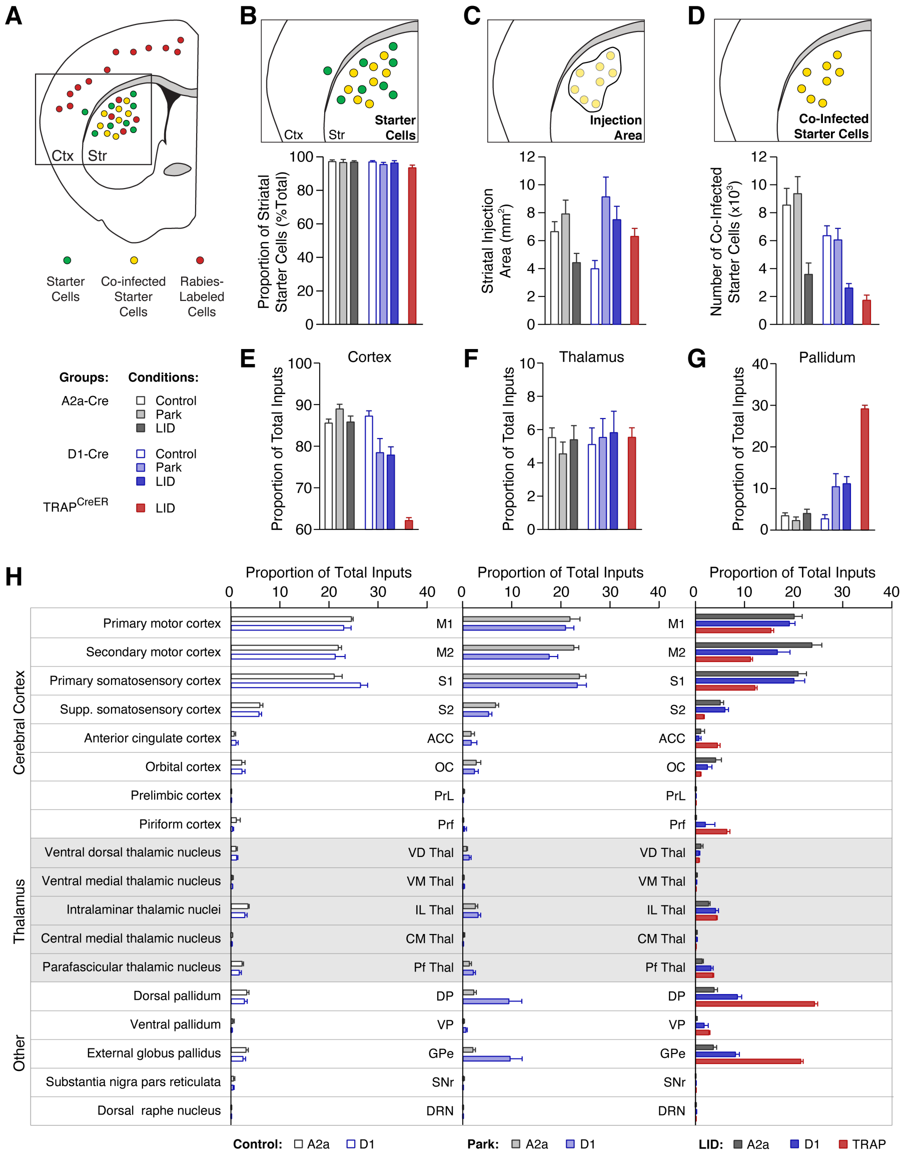


**Figure S2. Monosynaptic Rabies Tracing onto Indirect Pathway, Direct Pathway, and TRAPed Striatal Neurons. Related to Fig 3.**

A dual viral, Cre-dependent strategy was used to label monosynaptic inputs onto direct pathway, indirect pathway, and TRAPed striatal neurons. **(A)** Top: Coronal schematic of starter (green), rabies-labeled (red), and co-infected starter (yellow) cells. Bottom: Description of the groups and treatment conditions. **(B-D)** Top: coronal schematic showing quantification approach. Bottom: quantification of injection site. (B) The proportion of striatal starter cells (sTpEpB positive) compared to all brain-wide starter cells. (C) Striatal injection area, quantified by the extent of co-infected cells in the striatum. (D) The total number of co-infected striatal neurons. **(E-H)** The proportion of all extra-striatal rabies labeled cell bodies identified in the cortex (E), thalamus (F), and external globus pallidus (G), including quantification of specific sub-regions within these structures (H). A2a: Control, N=6, Park, N=4, LID, N=4; D1: Control, N=9, Park, N=10, LID, N=6; TRAP: LID, N=6. Data presented as mean ± SEM. N= animals.

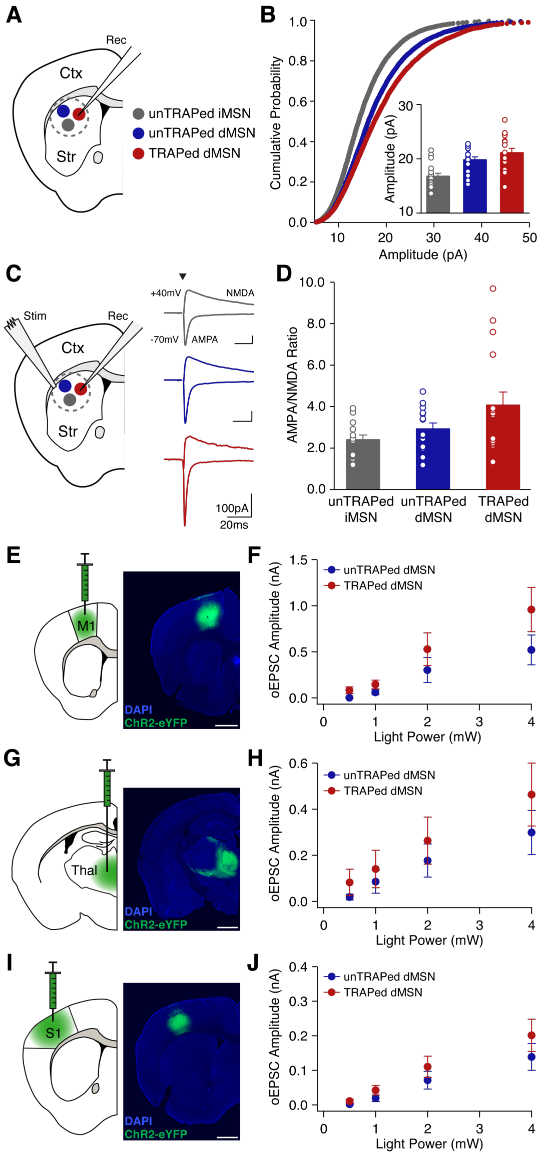


**Figure S3. Increased Motor Cortical and Thalamic Excitatory Transmission onto TRAPed dMSNs. Related to Figure 4.**

Excitatory inputs to striatal dMSNs were compared in *ex vivo* brain slices from the dorsolateral striatum of FosTRAPxAi14xD2-GFP (FAD) mice. **(A)** UnTRAPed iMSNs (gray), unTRAPed dMSNs (blue), and TRAPed dMSNs (red) were targeted for voltage-clamp recordings of miniature excitatory postsynaptic currents (mEPSCs). **(B)** Cumulative probability of mEPSC amplitudes. Inset: average mEPSC amplitude (unTRAPed iMSNs: n=21, N=8; unTRAPed dMSNs: n=20, N=7; TRAPed dMSNs: n=20, N=8). **(C)** (Left) UnTRAPed iMSNs (gray), unTRAPed dMSNs (blue), and TRAPed dMSNs (red) were targeted for voltage-clamp recordings of electrically evoked EPSCs. (Right) Representative EPSCs recorded at holding potentials of -70mV and +40mV to measure AMPA- and NMDA-mediated currents, respectively. Arrowheads denote timing of the electrical stimulus. **(D)** AMPA/NMDA ratio in unTRAPed iMSNs (n=14, N=7), unTRAPed dMSNs (n=16, N=9), and TRAPed dMSNs (n=16, N=8). N=mice, n=cells. **(E-J)** Excitatory inputs onto paired TRAPed and unTRAPed dMSNs were compared using an optogenetic approach. **(E,F)** Optical activation of primary motor cortical inputs (M1, n=19, N=4). **(G,H)** Optical activation of thalamic inputs (Thal, n=15, N=5). **(I,J)** Optical activation of primary somatosensory cortical inputs (S1, n=13, N=7). **(E,G,I)** Schematic diagrams (left) and histological sections (right) showing expression of Synapsin-ChR2-eYFP. Scale bar = 1 mm. **(F,H,J)** Average oEPSC amplitude versus light power for unTRAPed (blue) and TRAPed (red) dMSNs. n=pairs, N=mice. Data presented as mean ± SEM.

**EXPERIMENTAL MODEL AND SUBJECT DETAILS**
**Animals**

We used 3-9 month old C57Bl/6 mice of either sex. Hemizygous FosTRAP mice (Liqun Luo, Stanford) were bred to either wild-type C57Bl/6 mice (WT, Jackson Labs) or homozygous Ai14 mice (Jackson Labs) to yield FosTRAP or FosTRAP;Ai14 mice. Hemizygous D2-GFP mice (Gong et al., 2003) were bred against WT mice to produce D2-GFP animals. For slice electrophysiology experiments, hemizygous FosTRAP;Ai14 mice were bred to hemizygous D2-GFP mice to yield FosTRAP;Ai14;D2-GFP mice. For rabies tracing experiments, Drd1a (line EY217) and Adora2a (line KG139) BAC-Cre mice from the GENSAT Project were used (Gerfen et al., 2013). Animals were housed 1-5 per cage on a 12-hour light/dark cycle with *ad libitum* access to rodent chow and water. All behavioral manipulations were performed during the light phase. We complied with local and national ethical and legal regulations regarding the use of mice in research. All experimental protocols were approved by the UC San Francisco Institutional Animal Care and Use Committee.

### METHOD DETAILS

#### Key Resource Table

| Reagent or Resource | Source | Identifier |
|---|---|---|
| **Antibodies** | | |
| Rabbit anti-TH | Pel-Freez Biologicals | Cat# P40101-150, RRID: AB_2617184 |
| Chicken anti-TH | Millipore | Cat# AB9702, RRID: AB_570923, Lot #2586825 |
| Chicken anti-GFP | Abcam | Cat# ab13970; RRID: AB_300798 |
| Alexa Fluor 488- Donkey Anti-Rabbit IgG | Jackson ImmunoResearch Labs | Cat# 711-546-152, RRID: AB_2340619, Lot #128227 |
| Alexa Fluor568- Donkey anti-Rabbit IgG | Invitrogen | Cat# A10042; RRID: AB_2534017 |
| Alexa Fluor 647- Donkey Anti-Rabbit IgG | Jackson ImmunoResearch Labs | Cat# 711-606-152, RRID: AB_2340625, Lot #118661 |
| Alexa Fluor488- Donkey anti-Chicken IgY (IgG) | Jackson ImmunoResearch | Cat# 703-545-155; RRID: AB2340375 |
| Alexa Fluor647- Donkey anti-Chicken IgY (IgG) | Jackson ImmunoResearch | Cat# 703-606-155; RRID: AB_2340380 |
| **Bacterial and Virus Strains** | | |
| AAV5-EF1a-DIO-eYFP | UNC Vector Core | Lot #AV4310g |
| AAV5-EF1a-DIO-hCHR2(H134R)-eYFP-wpre-hGH | Penn Vector Core | AV-5-20298P, Lot #CS0384 |
| rAAV1/synp-DIO-sTbEpB-GFP | UNC Vector Core | Lot AV6118CD |
| EnvA G-deleted Rabies-mcherry | Salk Viral Vector Core | |
| **Chemicals, Peptides, and Recombinant Proteins** | | |
| Picrotoxin | Sigma-Aldrich | P1675 |
| Lidocaine N-ethyl chloride | Sigma-Aldrich | L1663 |
| 6-Hydroxydopamine hydrobromide | Sigma-Aldrich | H116 |
| Potassium methanesulfonate | Sigma-Aldrich | 83000 |
| Guanosine 5'-triphosphate sodium salt hydrate | Sigma-Aldrich | G8877 |
| Adenosine 5'-triphosphate magnesium salt | Sigma-Aldrich | A9187 |
| Desipramine hydrochloride | Sigma-Aldrich | D3900 |
| Benserazide hydrochloride | Sigma-Aldrich | B7283 |
| SKF 81297 hydrobromide | Tocris Bioscience | 1447 |
| 3,4-Dihydroxy-L-phenylalanine | Sigma-Aldrich | D9628 |
| Cesium methanesulfonate | Sigma-Aldrich | C1426 |
| Triton X-100 | Sigma-Aldrich | T8787 |
| Biocytin | Sigma-Aldrich | B4261 |
| 4-Hydroxytamoxifen | Sigma-Aldrich | H6278 |
| Tetrodotoxin | Abcam | ab120054 |
| **Critical Commercial Assays** | | |
| VECTASHIELD Antifade Mounting Medium | Vector Laboratories | Cat# H-1000, RRID: AB_2336789 |
| **Experimental Models: Organisms/Strains** | | |
| Mouse: WT: C57BL/6J | The Jackson Laboratory | RRID: IMSR_JAX:000664 |
| Mouse: B6.129(Cg)-Fostm1.1(cre/ERT2)Luo/J | The Jackson Laboratory | RRID: IMSR_JAX:021882 |
| Mouse: STOCK Tg(Drd2-EGFP)S118Gsat/Mmnc Mus musculus | MMRRC | RRID: MMRRC_000230-UNC |
| Mouse: B6.Cg-Gt(ROSA)26Sortm14(CAG-tdTomato)Hze/J | The Jackson Laboratory | RRID: IMSR_JAX:007914 |
| Mouse: B6.FVB(Cg)-Tg(Drd1-cre)EY217Gsat/Mmucd | MMRRC | RRID: MMRRC_034258-UCD |
| Mouse: B6.FVB(Cg)-Tg(Adora2a-cre)KG139Gsat/Mmucd | MMRRC | RRID: MMRRC:036158-UCD |
| **Software and Algorithms** | | |
| Igor Pro | Wavemetrics | http://www.wavemetrics.com/products/igorpro/igorpro.htm; RRID: SCR_000325 |
| MATLAB R2015a | MathWorks | https://www.mathworks.com/products/matlab/; RRID: SCR_001622 |
| ImageJ | NIH | https://imagej.nih.gov/ij/; RRID: SCR_003070 |
| EthoVision XT | Noldus | http://www.noldus.com/animal-behavior-research/products/ethovision-xt; RRID: SCR_000441 |
| Adobe Illustrator CS5 | Adobe | https://www.adobe.com/products/illustrator.html; RRID: SCR_014198 |
| MAP Software | Plexon | http://plexon.com/products/map-software; RRID: SCR_003170 |
| Offline Sorter | Plexon | http://plexon.com/products/offline-sorter; RRID: SCR_000012 |
| NeuroExplorer | NeuroExplorer | http://www.neuroexplorer.com/; RRID: SCR_001818 |
| mafPC | Xu-Friedman Lab | https://www.xufriedman.org/mafpc; |
| Axon MultiClamp Commander Software | Axon | http://mdc.custhelp.com/app/answers/detail/a_id/18877/~/axon%E2%84%A2-multiclamp%E2%84%A2-commander-software-download-page |
| NIS-Elements | Nikon | https://www.microscope.healthcare.nikon.com/products/software; RRID:SCR_014329 |
| NeuroInfo | MBF Biosciences | https://www.mbfbioscience.com/ |
| **Other** | | |
| 32-channel fixed optrode array | Innovative Neurophysiology | http://www.inphysiology.com/optogenetic-applications/ |
| 200 µm Core TECS-Clad Multimode Optical Fiber, 0.39 NA | Thorlabs | Cat# FT200UMT |
| 1.25 mm Multimode LC/PC Ceramic Ferrule, 230 µm Bore Size | Thorlabs | Cat# CFLC230-10 |
| Ceramic Split Mating Sleeve for 1.25 mm (LC/PC) Ferrules | Thorlabs | Cat# ADAL1 |
| 150mW DPSS 473nm Blue Laser | Shanghai Laser & Optics Century | BL473T8-150 + ADR-800A |
| 200 µm Core, 0.39 NA FC/PC to Ø1.25 mm Ferrule Patch Cable, 1 m Long | Thorlabs | Cat# M83L01 |
| 1x1 Fiber-optic Rotary Joint | Doric Lenses | Cat# FRJ_1x1_FC-FC |
| 1x2 Fiber-optic Rotary Joint | Doric Lenses | Cat# FRJ_1x2i_FC-2FC_0.22 |
| 10-Channel Slip Ring Electrical Commutator | Dragonfly | Model SL-10-C; https://campdeninstruments.com/products/10-ch-slip-ring-commutator-30g |
| Master-8 | A.M.P.I. | http://www.ampi.co.il/master8cp.html |
| M-Series (M60) Amplifier System for Multiplexing | Triangle BioSystems | http://www.trianglebiosystems.com/m-series-systems.html |
| Single Channel Temperature Controller | Warner Instruments | TC-324C |
| MINIPULS 3 Peristaltic Pumps | Gilson | F155008 |
| X-Cite 120LED Boost | Excelitas | 010-00326R |
| ITC-18 16-bit Multi-Channel Data Acquisition Interface | Heka | ITC-18 |
| MultiClamp 700B Microelectrode Amplifier | Molecular Devices | |

#### Surgical Procedures

A detailed protocol for stereotaxic surgery can be found at dx.doi.org/10.17504/protocols.io.n2bvj6qynlk5/v1. Briefly, all surgical procedures were performed at 3-6 months of age. Anesthesia was induced with intraperitoneal (IP) injection ketamine/xylazine and maintained with 0.5%-1.0% inhaled isoflurane. Mice were placed in a stereotaxic frame and a mounted drill was used to create holes over the left medial forebrain bundle (MFB), the left dorsolateral striatum (DLS), primary motor cortex (M1), primary somatosensory cortex (S1), or thalamus (Thal). To render mice parkinsonian, the left MFB (-1.0 AP, +1.0 ML, -4.9 mm DV) was injected using a 33-gauge needle with 1-1.5 µL per site of 6-Hydroxydopamine (6-OHDA)-

bromide. In some experiments, AAV5-DIO-ChR2-eYFP (UPenn Vector Core, 1-1.5 μL, diluted 1:3 in NS) was injected in the left DLS (+0.8 AP, +2.3 ML, -2.5 mm DV). For Cre-dependent rabies tracing experiments, 300 nL helper virus, rAAV1/synp-DIO-sTbEpB-GFP (UNC Vector Core, lot AV6118CD) was injected into two left DLS sites (-0.8 AP, -2.4 ML, -2.5 DV). Two weeks after helper virus injection, 300nL modified EnvA G-deleted Rabies-mcherry (Salk Viral Vector Core) virus was also injected into the same DLS site at 100 nL/min. For input-specific circuit mapping onto TRAPed cells, 250-300nL of AAV5-hSyn-hChR2(H134R)-eYFP (UNC Vector Core) was injected into M1 (+1.2 AP, -1.6 ML, -0.7 mm DV), S1 (+0.95 AP, -2.9 ML, -0.75 mm DV), or thalamus (-2.3 AP, -0.6 ML, +4.0 mm DV) in FosTRAP;WT mice. 6-OHDA and virus were injected at a rate of 0.10 μL/min, after which the injection cannula was left in place for 10-15 minutes prior to being withdrawn and the scalp being sutured.

In preparation for *in vivo* single-unit recordings, FosTRAP;Ai14 mice were injected with 6-OHDA and DIO-ChR2, as described above, and optrode arrays were implanted in a second surgical procedure. After the scalp was reopened, a large craniotomy (1.5 x 1 mm) was created over the left DLS, and two small holes were drilled in the right frontal and right posterior parietal areas for placement of a skull screw (Fine Scientific Tools, FST) and ground wire, respectively. A fixed multichannel electrode array (32 Tungsten microwires, Innovative Neurophysiology) coupled to a 200 μm optical fiber (Thorlabs) was slowly lowered through the craniotomy into the DLS. The final location of the electrode tips was targeted 100-200 μm above the previous DIO-ChR2 injection (-2.3-2.4 mm DV). The array was covered and secured into place with dental cement (Metabond) and acrylic (Ortho-Jet).

All animals were given buprenorphine (IP, 0.05 mg/kg) and ketoprofen (subcutaneous injection, 5 mg/kg) for postoperative analgesia. Parkinsonian animals were monitored closely for 1 week following surgery: mouse cages were kept on a heating pad; animals received daily saline injections and were fed nutritional supplements (Diet-Gel Recovery Packs and forage/trail mix).

**Behavior**

Postoperatively, parkinsonian mice were monitored in the open field 1-2 times per week for 10 minutes per session. A detailed protocol can be found at dx.doi.org/10.17504/protocols.io.n2bvj6qynlk5/v1. Briefly, all mice were habituated to the open field (clear acrylic cylinders, 25 cm diameter) for 30 minutes 1-2 days prior to behavioral sessions. The mice were monitored via two cameras, one directly above (to capture overall movement) and one in front of the chamber (to capture fine motor behaviors). Video-tracking software (Noldus Ethovision) was used to quantify locomotor activity, including rotations (90° contralateral or ipsilateral turns), distance traveled, and velocity. After a three-week baseline period, daily injections of levodopa commenced. Levodopa-induced dyskinesia (LID) was scored during weekly sessions in which mice were injected, then placed in a clean, clear cage for visualization. For regular weekly dyskinesia scoring, 1-2 blinded experimenters rated AIMs (for details see Statistical Procedures below). For *in vivo* electrophysiology experiments, rotations and AIMs were quantified in one-minute bins, with dyskinesia being scored every other minute.

**Pharmacology**

6-OHDA (Sigma Aldrich) for MFB dopamine depletions was prepared at 5 μg/μL in normal saline solution. Levodopa (Sigma Aldrich) was administered with benserazide (Sigma Aldrich) and prepared in normal saline solution. Levodopa (5-10 mg/kg) was given via IP injection 5-7 days per week over the course of the experiment. Initially, on the $7^{th}$ day of levodopa treatment for FosTRAP;WT, FosTRAP;Ai14, and FosTRAP;Ai14;D2-GFP mice were given 4-hydroxytamoxifen (4-OHT, 50 mg/kg in Chen oil, IP) exactly one hour post-levodopa injection, to capture dyskinesia-associated neurons (Figure 1A). 4-OHT was prepared as previously described (Guenthner et al., 2013). Briefly, to prepare a 20 mg/mL stock in ethanol of 4-OHT, 4-OHT was added to 200 proof ethanol, vortexed, and placed on a horizontal shaker at 37° C for 30 minutes or until the 4-OHT dissolved. The stock solution was kept covered in foil to minimize light exposure. Next, to prepare a 10 mg/mL working solution in oil, the 4-OHT/ethanol mixture was combined with Chen Oil (a mixture of 4 parts sunflower seed oil and 1 part castor oil) and placed into 1.5 mL Eppendorf tubes. The tubes were vigorously mixed, wrapped in foil, and left on a nutator for 45 min at room temperature, vortexed and shaken periodically. The tubes were then placed in a speed-vac for 2-3 hours to evaporate the ethanol. If necessary, the final volume was adjusted with Chen Oil to 1 mL to reach a final concentration of 10 mg/mL. Both levodopa and 4-OHT were injected in a quiet, familiar environment, and animals were returned to their home cages, to minimize additional stimuli. Daily levodopa injections continued for 2-6 weeks to allow expression of Cre-dependent constructs. For *ex vivo* experiments, picrotoxin (Sigma) was dissolved in warm water to prepare a 5 mM stock solution, which was subsequently diluted in ACSF for a final concentration of 50 µM. Tetrodotoxin (TTX, Abcam) was dissolved in water at a stock concentration of 1 mM and added to ACSF for a final concentration of 1 µM. SKF 81298 (Tocris) was dissolved in water at a concentration of 1mM and added to ACSF for a final concentration of 5 µM. For all *ex vivo* experiments, biocytin (1-2.5 mg/mL) was included in the internal solution for post-hoc confirmation of the presence or absence of Ai14 and D2-GFP.

#### ***In Vivo* Electrophysiology**

A detailed protocol for in vivo electrophysiology, including optrode array fabrication and data acquisition, can be found at dx.doi.org/10.17504/protocols.io.5jyl89w69v2w/v1. Two weeks after optrode array implantation, mice were habituated to tethering and the recording chamber for 1-2 days. After habituation, experimental sessions occurred 3-5 times per week for 2-6 weeks. During each session, electrical signals (single-unit and LFP data from each of 32 channels) were collected using a multiplexed 32 channel headstage (Triangle Biosystems), an electrical commutator equipped with a fluid bore (Dragonfly), filtered, amplified, and recorded on a MAP system, using RASPUTIN 2.4 HLK3 acquisition software (Plexon). Spike waveforms were filtered at 154–8800 Hz and digitized at 40 kHz. The experimenter manually set a gain and threshold for storage of electrical events.

During recording sessions, after a baseline period of 30 minutes in the parkinsonian state, levodopa (5-10 mg/kg) was injected IP. After a period of 2-3 hours of recording spontaneous activity in the open field, an optogenetic cell identification protocol was applied (Girasole et al., 2018) consisting of 100 msec blue light pulses, given at 1 Hz. At each of 4 light powers (0.5, 1, 2, and 4 mW), 1000 light pulses were delivered via a lightweight patch cable (Doric Lenses) connected to a blue laser (Shanghai Laser and Optics Century), via an optical commutator (Doric Lenses), and controlled by TTL pulses from a behavioral monitoring system (Noldus Ethovision).

Single-units were identified offline by manual sorting using Offline Sorter 3.3.5 (Plexon) and principal components analysis (PCA). Clusters were considered to represent a single unit if (1) the unit's waveforms were statistically different from multiunit activity and any other single-units on the same wire, in 3D PCA space, (2) no interspike interval <1 msec was observed. Single-units were then classified as putative medium spiny neurons (MSNs) or interneurons (INs) as previously described (Berke et al., 2004; Gage et al., 2010; Harris et al., 2000; Barnes et al., 2005) using features of the spike waveform (peak to valley and peak width), as well as inter-spike interval distribution.

After single-units had been selected for further study, their firing activity was analyzed using NeuroExplorer 4.133 (Nex Technologies). To determine if a unit was optogenetically identified, a peristimulus time histogram was constructed around the onset of laser pulses. To be considered optogenetically identified, a unit had to fulfill 3 criteria: (1) the unit had to increase firing rate above the 99% confidence interval of the baseline within 15 msec of laser onset; (2) the unit's firing was above this threshold for at least 15 msec; (3) the unit's laser-activated waveforms were not statistically distinguishable from spontaneous waveforms.

#### *Ex vivo* Electrophysiology

A detailed protocol for slice electrophysiology can be found at dx.doi.org/10.17504/protocols.io.6qpvr67rpvmk/v1. Prior to terminal anesthesia and preparation of brain slices, animals (3-9 months) were co-injected with levodopa and benserazide (5-10 mg/kg and 2.5-5 mg/kg, respectively) to induce LID. After 30-45 minutes in the dyskinetic state, mice were deeply anesthetized with an IP ketamine-xylazine injection, transcardially perfused with ice-cold glycerol-based slicing solution, decapitated, and the brain was removed. Glycerol-based slicing solution contained (in mM): 250 glycerol, 2.5 KCl, 1.2 $NaH_2PO_4$, 10 HEPES, 21 $NaHCO_3$, 5 glucose, 2 $MgCl_2$, 2 $CaCl_2$. The brain was mounted on a submerged chuck, and sequential 275 µm coronal slices were cut on a vibrating microtome (Leica), transferred to a chamber of warm (34° C) carbogenated ACSF containing (in mM) 125 NaCl, 26 $NaHCO_3$, 2.5 KCl, 1 $MgCl_2$, 2 $CaCl_2$, 1.25 $NaH_2PO_4$, 12.5 glucose for 30-60 minutes, then stored in carbogenated ACSF at room temperature. Each slice was then submerged in a chamber superfused with carbogenated ACSF at 31-33°C for recordings.

Striatal medium spiny neurons were targeted for recordings using differential interference contrast (DIC) optics in FosTRAP;Ai14;D2GFP mice on a Olympus BX 51 WIF microscope. In FosTRAP;Ai14;D2GFP mice, TRAPed neurons were identified by their tdTomato-positive somata and D2-positive neurons were identified by GFP fluorescence. Fluorescence-negative neurons with GABAergic interneuron physiological properties (membrane tau decay <0.8 ms for both fast-spiking and persistent low-threshold spiking subtypes; input resistance >500 MΩ in persistent low-threshold spiking subtype) were excluded from the analysis.

Neurons were patched in whole-cell voltage-clamp configurations using borosilicate glass electrodes (3-5 MΩ) filled with cesium-based (voltage-clamp) or potassium methanesulfonate-based (current-clamp) internal solution. Cesium based solution containing (in mM) respectively: 120 $CsMeSO_3$, 15 CsCl, 8 NaCl, 0.5 EGTA, 10 HEPES, 2 MgATP, 0.3 NaGTP, 5 QX-314, pH 7.3. Potassium based solution containing (in mM): 130 $KMeSO_3$, 10 NaCl, 2 $MgCl_2$, 0.16 $CaCl_2$, 0.5 EGTA, 10 HEPES, 2 MgATP, 0.3 NaGTP, pH 7.3. Picrotoxin (50 µM) was added to the external solution to block synaptic currents mediated by $GABA_A$ receptors. Drugs were prepared as stock solutions and added to the ACSF to yield the final concentration.

Whole-cell recordings were made using a MultiClamp 700B amplifier (Molecular Devices) and ITC-18 A/D board (HEKA). Data was acquired using Igor Pro 6.0 software (Wavemetrics) and custom acquisition routines (mafPC, courtesy of M. A. Xu-Friedman). Both voltage clamp and current-clamp recordings were filtered at 2 kHz and digitized at 10 kHz. All recorded neurons exhibited electrophysiological characteristics of medium spiny neurons. All synaptic currents were recorded with a cesium-based internal and monitored at a holding potential of -70 mV. Series resistance and leak currents were monitored continuously. Miniature EPSCs were recorded at – 70 mV in 1 μM TTX and 50 μM picrotoxin. Evoked EPSCs onto medium spiny neurons were elicited in the presence of picrotoxin (50 μM) with a stimulus isolator (IsoFlex, AMPI) and a glass electrode placed dorsolateral to the recorded neuron, typically 100-200 μm away. Stimulus intensity was adjusted to yield EPSC amplitudes of approximately 400 pA with a stimulus duration of 300 μs. For evaluation of the paired pulse ratio, two stimuli were given at variable interstimulus intervals (ISIs; 25, 50, 100, 200, 500 ms) with a 20 sec intertrial interval. Paired-pulse ratio is defined as $EPSC_2/EPSC_1$. Five-eight repetitions at each ISI were averaged to yield the PPR for that ISI. For monitoring of EPSC amplitude over time, two pulses delivered with 50 ms interstimulus interval were given every 20 seconds. For AMPA/NMDA ratio experiments, one stimulus at -70 mV or +40 mV was given every 20 seconds, at 15-20 repetitions per holding potential. AMPA/NMDA ratios were calculated as the ratio of the magnitude of the EPSC at +40 mV at 50 ms following stimulation (NMDA) to the peak of the EPSC at -70 mV (AMPA). To evoke ChR2-mediated synaptic currents from M1, S1, or thalamus, excitatory currents were optically evoked using 2 ms pulses of 473 nm light at light powers of 0.5, 1, 2, and 4 mW and delivered by a TTL-controlled LED (Olympus) passed through a GFP filter (Chroma).

Current-clamp recordings were made to measure intrinsic properties of striatal neurons. The resting membrane potential ($V_m$) was measured as the average $V_m$ 5-10 minutes after break-in. A series of small negative current steps were delivered from rest to calculate the input resistance of each cell. Rheobase and other input-output properties were obtained by giving a series of square-wave current steps, ranging from 100 pA to 600 pA, in 100 pA increments, with a 10 sec interstimulus interval. Drugs, such as SKF-81297, were applied after achieving a stable baseline 5-10 minutes after break-in. Changes in intrinsic properties due to SKF were assessed 10-15 minutes after drug wash-in.

**Monosynaptic Rabies Tracing**

D1-Cre and A2a-Cre mice were used to perform monosynaptic retrograde tracing onto direct and indirect pathway neurons, respectively. Groups of healthy (non-depleted), parkinsonian, and parkinsonian/levodopa-treated mice were used within each genotype. Mice were rendered parkinsonian as described above. Four weeks after dopamine depletion, animals received daily injections of levodopa. Parkinsonian mice (one week into daily levodopa injections) or untreated healthy mice, were then anesthetized and a Cre-dependent helper virus (AAV-DIO-sTpEpB-GFP) was stereotaxically injected into the left DLS (ipsilateral to the depletion in parkinsonian mice). The helper virus (AAV-DIO-sTpEpB-GFP) expresses the EnvA receptor (TVA) and rabies glycoprotein necessary for rabies infection and replication in a cell-type specific manner, termed "starter cells," which are labeled with the green fluorophore GFP. After animals recovered for two weeks, they were anesthetized and a replication-incompetent form of the rabies virus (EnvA-G-deleted-rabies-mCherry) was stereotaxically injected into the DLS using the same coordinates. The rabies virus will then infect a subset of starter cells (co-infected) and travel retrogradely one synapse, expressing the red fluorophore mCherry in infected cells. Once the rabies virus infects a presynaptic neuron, uninfected with the helper virus, it will no longer be capable of replication and/or retrograde synaptic infection. Rabies injections were performed in an approved Biosafety Level 2 (BSL-2) surgical suite. Animals were then allowed to recover for ten days, at which point they were terminally anesthetized with ketamine/xylazine (200/40 mg/kg I.P.), transcardially perfused with 4% paraformaldyde (PFA), and the brain dissected from the skull. The brain was post-fixed overnight in 4% PFA and then placed in 30% sucrose at 4°C.

Parkinsonian, levodopa-treated FosTRAP;WT mice were prepared in a similar fashion as D1-Cre and A2a-Cre mice, with some alterations made to the experimental timeline to accommodate helper virus expression using the conditional Cre (CreER) in the FosTRAP line. In FosTRAP mice, helper virus was injected in the left DLS at the same time as the initial dopamine depletion. Three weeks after dopamine depletion, FosTRAP mice began daily levodopa injections. After one week of daily levodopa injections, as above, FosTRAP mice were injected with levodopa followed by an injection of 4-OHT, allowing for recombination and expression of the helper virus. Two weeks later, FosTRAP mice were anesthetized and the modified rabies virus was injected using the same procedures described above. The remainder of the experimental timeline was similar to that for D1-Cre and A2a-Cre mice as described above.

Fixed brains, stored in sucrose, were then sent to Dr. Charles Gerfen at the National Institutes of Mental Health (NIMH) for sectioning, mounting, imaging, and analysis using published methods (Eastwood et al., 2019). Briefly, brains were sectioned coronally at 50 μm using a freezing microtome. Sections were processed for fluorescent immunohistochemical localization of GFP labeling of rabies starter cells, RFP labeling of transsynaptically transported rabies, and tyrosine hydroxylase to label the nigrostriatal dopamine system. Slices were then imaged using a Zeiss microscope equipped with a z axis drive, imaging each fluorophore. The imaged coronal sections were reconstructed into a whole brain volume, labeled cells detected using a modified Laplacian of Gaussian algorithm and then registered to the Allen Common Coordinate mouse atlas framework using NeuroInfo software (MBF Biosciences, Williston, VT).

**Histology & Microscopy**

After rabies tracing or behavioral experiments, mice were deeply anesthetized with IP ketamine-xylazine and transcardially perfused with 4% paraformaldehyde in PBS. Following *in vivo* electrophysiology experiments, prior to perfusion, electrode array location was marked by electrolytic lesioning. After deep anesthesia, the implant was connected to a solid state, direct current (DC) Lesion Maker (Ugo Basile). A current of 100 μA was passed through each microwire for 5 seconds. After perfusion, the brain was dissected from the skull and post-fixed overnight in 4% paraformaldehyde, then placed in 30% sucrose at 4°C for cryoprotection. The brain was then cut into 35 µm coronal or sagittal sections on a freezing microtome (Leica) and then mounted in Vectashield Mounting Medium onto glass slides for imaging. For immunohistochemistry, the tissue was blocked with 3% normal donkey serum (NDS) and permeabilized with 0.1% Triton X-100 for 2 hours at room temperature on a shaker. Primary antibodies were added to 3% NDS and incubated overnight at 4° C on a shaker. Primary antibodies used: Rabbit anti-TH (Pel-Freez, 1:1000), Chicken anti-TH (Sigma, 1:1000), and Chicken anti-GFP (1:500). Slices were then incubated in secondary antibodies (donkey anti-rabbit or chicken Alexafluor 488, 593, or 647, 1:500, JacksonImmuno Research) for 2-4 hours at 4°C on a shaker, washed, and mounted onto slides for imaging. 4 or 10x images were acquired on a Nikon 6D conventional widefield microscope.

For slice electrophysiology experiments in which the internal solution contained biocytin, slices were subsectioned at 50 µm and washed in PBS. Slices were blocked for 2 hours at room temperature on a shaker in a 5% NDS and 0.3% Tween-20 PBS-based solution. Primary antibodies were the same as described above. Slices were then incubated in secondary antibodies (donkey anti-rabbit or chicken Alexafluor 488, 593, or 647, 1:500, JacksonImmuno Research and Streptavidin Alexa 350, 3:500, Sigma) for 6-12 hours at 4°C on a shaker, washed, and mounted onto slides for imaging. Images were acquired on a Nikon 6D conventional widefield or Nikon Spinning Disk confocal microscope with a 40x objective microscope. Exposure times were matched between images of the same type. Post-hoc confirmation of cellular identify of a subset of recovered biocytin-filled, recorded cells revealed that online identification by experimenter using fluorescence intensity was >90% in FosTRAP;Ai14;D2-GFP mice for TRAPed dMSNs and unTRAPed dMSNs and iMSNs. The rate of positive identification of TRAPed and unTRAPed dMSNs in FosTRAP:Ai14;D2-GFP mice injected with ChR2-eYFP in M1, S1, or Thal was also > 90%, however, due to the overlap of YFP and GFP emission spectra, our positive rates of identification of unTRAPed iMSNs was reduced to ~60%, leading to the exclusion of unTRAPed iMSNs from these experiments (Figure 4D-N, Figure SG-O).

### QUANTIFICATION AND STATISTICAL ANALYSIS

#### Statistics

Details can be found in the Statistical Table. All data are presented as the mean ± SEM, with N referring to the number of animals and n to the number of cells. Goal sample sizes for physiological studies were chosen using a power calculation, with a two-sided alpha of 0.05, power of 0.9, and the statistical tests listed below under each section. For behavioral and *in vivo* physiology assays, goal sample size was driven by the lowest-yield component of the experiments (optogenetically-labeled single-unit recordings of TRAPed neurons, goal n=10). The power calculation relied on estimates of effect size from DYSK vs ON dMSN subtypes (Ryan et al., 2018). For *ex vivo* physiology assays, the power calculation relied on previously acquired data in the lab, in another mouse model of dyskinesia, to estimate average, standard deviation, and effect size. Power calculations yielded n=15 cells/cell type per group for mEPSCs, and n=10 cells/group for evoked EPSC analyses. For *ex vivo* pharmacology, we again used previously acquired data in the lab, as well as published data on the effects of D1 agonists (Planert et al., 2013) to calculate the sample size, which yielded n=7 cells per group. For all *ex vivo* electrophysiology, regardless of the minimum n (cells), we had a goal of N>4 (mice). For modified rabies tracing, there were few precedents in the literature to inform a power calculation, so we chose a goal sample size of N=5 mice per group, per cell type.

| Key Experiments | Figure | Type of Comparison | Statistical Test | n (units/cells) | N (animals) | Comparison values (± SEM) & p value | Planned Sample Size (from power calculation) |
|---|---|---|---|---|---|---|---|
| **Baseline firing rate of MSNs in Park state, in vivo** | 1k | Between-group | MWU | 8 (opto labeled TRAP dMSN), 117 (dMSN) | 6 (TRAP), 14 (dMSN) | TRAP dMSNs: 0.14 ± 0.07 Hz; All dMSNs: 0.14 ± 0.03 Hz; p=0.773 | n = 10 cells |
| **Change in firing rate of MSNs in LID state, in vivo** | 1k | Between-group | MWU | 8 (opto labeled TRAP dMSN), 117 (dMSN) | 6 (TRAP), 14 (dMSN) | TRAP dMSNs: 5.92 ± 1.92 Hz, All dMSNs: 2.62 ± 0.32 Hz; p=0.033 | n = 10 cells |
| **Correlation of MSN firing rate to dyskinesia severity** | 1o, S1j | Between-group | MWU | 8 (opto labeled TRAP dMSN), 117 (dMSN) | 6 (TRAP), 14 (dMSN) | TRAP dMSNs: R2 = 0.39 ± 0.06; All dMSNs: R2 = 0.24 ± 0.02; p=0.033 | n = 10 cells |
| **Evoked firing rate of MSNs in Park state, ex vivo** | 2f, Table 1 | Between-group | rmANOVA | 22 (TRAP dMSN), 17 (un T dMSN), 17 (unT iMSN) | 14 (TRAP dMSN), 13 (un T dMSN), 11 (unT iMSN) | F(10,230)=2.598, p=0.005, unT dMSN vs. unT iMSN: p=0.197, TRAP dMSN vs. unT dMSN: p=0.999, TRAP dMSN vs unT iMSN: p=0.187 | n=15 cells/group |
| **Changes in evoked firing rate of MSNs after SKF-81297, ex vivo** | 2g, Table 2 | Within-cell | rmANOVA | 14 (TRAP dMSN), 11 (un T dMSN), 9 (unT iMSN) | 10 (TRAP dMSN), 9 (un T dMSN), 6 (unT iMSN) | unT iMSN: F(5,80)=0.362, p=0.874, unT dMSN: F(5,110)=0.370, p=0.839, TRAP dMSN: F(5,130)=2.375, p=0.0425 | n=10 cells/group |
| **Total number of presynaptic inputs onto MSNs (control)** | 3h | Between-group | MWU | n/a | 6 (A2a), 9 (D1) | Control: iMSN vs dMSN, p=0.114 | N = 10 mice/group |
| **Number of cortical inputs onto iMSNs (all groups)** | 3i | Between-group | KW | n/a | Control: 6 (A2a), Park: 4 (A2a), LID: 4 (A2a) | χ2(2) 8.881, p=0.012 | N = 5 mice/group |
| **Number of cortical inputs onto iMSNs (control vs. Park)** | 3i | Between-group | HSD | n/a | Control: 6 (A2a), Park: 4 (A2a) | Control vs Park, p=0.0498 | N = 5 mice/group |
| **Number of cortical inputs onto iMSNs (Park vs. LID)** | 3i | Between-group | HSD | n/a | Park: 4 (A2a), LID: 4 (A2a) | Park vs LID, p=0.970 | N = 5 mice/group |
| **Number of cortical inputs onto iMSNs (control vs. LID)** | 3i | Between-group | HSD | n/a | Park: 4 (A2a), LID: 4 (A2a) | Healthy vs LID, p=0.024 | N = 5 mice/group |
| **Number of thalamic inputs onto iMSNs (all groups)** | 3j | Between-group | KW | n/a | Control: 6 (A2a), Park: 4 (A2a), LID: 4 (A2a) | χ2(2) 8.214, p=0.017 | N = 5 mice/group |
| **Number of thalamic inputs onto iMSNs (control vs. Park)** | 3j | Between-group | HSD | n/a | Control: 6 (A2a), Park: 4 (A2a) | Control vs Park, p=0.184 | N = 5 mice/group |
| **Number of thalamic inputs onto iMSNs (Park vs. LID)** | 3j | Between-group | HSD | n/a | Park: 4 (A2a), LID: 4 (A2a) | Park vs LID, p=0.343 | N = 5 mice/group |
| **Number of thalamic inputs onto iMSNs (control vs. LID)** | 3j | Between-group | HSD | n/a | Control: 6 (A2a), LID: 4 (A2a) | Control vs LID, p=0.0151 | N = 5 mice/group |
| **Number of GPe inputs onto iMSNs (all groups)** | 3k | Between-group | KW | n/a | Control: 6 (A2a), Park: 4 (A2a), LID: 4 (A2a) | χ2(2) 8.2571, p=0.016 | N = 5 mice/group |
| **Number of GPe inputs onto iMSNs (control vs. Park)** | 3k | Between-group | HSD | n/a | Control: 6 (A2a), Park: 4 (A2a) | Control vs Park, p=0.1714 | N = 5 mice/group |
| **Number of GPe inputs onto iMSNs (Park vs. LID)** | 3k | Between-group | HSD | n/a | Park: 4 (A2a), LID: 4 (A2a) | Park vs LID, p=0.0286 | N = 5 mice/group |
| **Number of GPe inputs onto iMSNs (control vs. LID)** | 3k | Between-group | HSD | n/a | Control: 6 (A2a), LID: 4 (A2a) | Control vs LID, p=0.019 | N = 5 mice/group |
| **Number of cortical inputs onto dMSNs (all groups)** | 3i | Between-group | KW | n/a | Control: 9 (D1), Park: 10 (D1), LID: 6 (D1) | χ2(2) 4.185, p=0.123 | N = 5 mice/group |
| **Number of thalamic inputs onto dMSNs (all groups)** | 3j | Between-group | KW | n/a | Control: 9 (D1), Park: 10 (D1), LID: 6 (D1) | χ2(2) 1.161, p=0.560 | N = 5 mice/group |
| **Number of Gpe inputs onto dMSNs (all groups)** | 3k | Between-group | KW | n/a | Control: 9 (D1), Park: 10 (D1), LID: 6 (D1) | χ2(2) 12.309, p=0.002 | N = 5 mice/group |
| **Number of GPe inputs onto dMSNs (control vs. Park)** | 3k | Between-group | HSD | n/a | Control: 9 (D1), Park: 10 (D1) | Control vs Park, p=0.0497 | N = 5 mice/group |
| **Number of GPe inputs onto dMSNs (Park vs. LID)** | 3k | Between-group | HSD | n/a | Park: 10 (D1), LID: 6 (D1) | Park vs LID, p=0.353 | N = 5 mice/group |
| **Number of GPe inputs onto dMSNs (control vs. LID)** | 3k | Between-group | HSD | n/a | Control: 9 (D1) , LID: 6 (D1) | Control vs LID, p<0.001 | N = 5 mice/group |
| **Total number of presynaptic inputs on dMSN (TRAP vs. LID)** | 3h | Between-group | MWU | n/a | LID: 6 (D1), LID: 6 (TRAP) | TRAP vs dMSN, p=0.132 | N = 5 mice/group |
| **Number of cortical inputs onto dMSNs (TRAP vs. LID)** | 3i | Between-group | MWU | n/a | LID: 6 (D1), LID: 6 (TRAP) | TRAP vs dMSN, p=0.009 | N = 5 mice/group |
| **Number of thalamic inputs onto dMSNs (TRAP vs. LID)** | 3j | Between-group | MWU | n/a | LID: 6 (D1), LID: 6 (TRAP) | TRAP vs dMSN, p=0.041 | N = 5 mice/group |
| **Number of GPe inputs onto dMSNs (TRAP vs. LID)** | 3k | Between-group | MWU | n/a | LID: 6 (D1), LID: 6 (TRAP) | TRAP vs dMSN, p=0.240 | N = 5 mice/group |
| **mEPSC amplitues of unTRAP dMSNs vs unTRAPed iMSNs** | 4c | Between-group | MWU | 20 (unT dMSN), 21 (unT iMSN) | 7 (un T dMSN), 8 (unT iMSN) | unT dMSN: 19.89 ± 0.47 pA, unT iMSN: 16.86 ± 0.54 pA, p<0.001 | n = 20 cells/group |
| **mEPSC amplitues of unTRAP dMSNs vs TRAPed dMSNs** | 4c | Between-group | MWU | 20 (TRAP dMSN), 20 (unT dMSN) | 8 (TRAP dMSN), 7 (unT dMSN) | TRAP dMSN: 21.22 ± 0.67 pA, unT dMSN: 19.89 ± 0.47 pA, p=0.102 | n = 20 cells/group |
| **mEPSC frequency of unTRAP dMSNs vs unTRAPed iMSNs** | 4d | Between-group | MWU | 20 (unT dMSN), 21 (unT iMSN) | 7 (unT dMSN), 8 (unT iMSN) | unT iMSN: 2.80 ± 0.36 Hz, unT dMSN: 3.87 ± 0.41 Hz, p=0.034 | n = 20 cells/group |
| **mEPSC frequency of unTRAP dMSNs vs TRAPed dMSNs** | 4d | Between-group | MWU | 20 (TRAP dMSN), 20 (unT dMSN) | 8 (TRAP dMSN), 7 (unT dMSN) | TRAP dMSN: 5.41 ± 0.47 Hz, unT dMSN: 3.87 ± 0.41 Hz, p=0.021 | n = 20 cells/group |
| **Evoked EPSCs, AMPA/NMDA ratio in dMSNs** | 4g | Between-group | MWU | 16 (TRAP dMSN), 16 (un T dMSN), 14 (unT iMSN) | 8 (TRAP dMSN), 9 (unT dMSN), 7 (unT iMSN) | TRAP dMSN: 4.07 ± 0.63, unT dMSN: 2.92 ± 0.28, p=0.376, RS | n=15 cells/group |
| **Evoked EPSCs, PPR** | 4i | Between-group | rmANOVA, HSD | 22 (TRAP dMSN), 18 (un T dMSN), 17 (unT iMSN) | 9 (TRAP dMSN), 9 (unT dMSN), 8 (unT iMSN) | F(10,270)=2.493, p=0.007, rmANOVA; unT dMSN vs unT iMSN, p=0.367, unT dMSN vs T dMSN, p=0.0186, unT iMSN vs T dMSN, p<0.001, Tukey | n=15 cells/group |
| **Optically evoked EPSCs, M1** | 4 | Between-group (sequential pairs) | WSR | 19 | 4 | T dMSN: 1.06 ± 0.25 nA, unT dMSN: 0.42 ± 0.14 nA, p=0.002, SR | n=15 pairs |
| **Optically evoked EPSCs, S1** | 4 | Between-group (sequential pairs) | WSR | 13 | 7 | T dMSN: 0.21 ± 0.05 nA, unT dMSN: 0.13 ± 0.04 nA, p=0.073, SR | n=15 pairs |
| **Optically evoked EPSCs, thalamus** | 4 | Between-group (sequential pairs) | WSR | 15 | 5 | T dMSN: 0.59 ± 0.18 nA, unT dMSN: 0.32 ± 0.11 nA, p=0.013, RS | n=15 pairs |

**Abbreviations:** WSR (Wilcoxon Signed Rank Test); MWU (Wilcoxon Rank Sum or Mann-Whitney U Test); rmANOVA (Repeated Measures Analysis of Variance); KW ( Kruskal-Wallis), HSD (Tukey's HSD)

### Behavior

Dyskinesia was quantified using a standard scoring method (Cenci and Lundblad, 2007), which takes into account abnormal involuntary movements (AIMs) in axial, limb, and orofacial (ALO) body segments. Briefly, dyskinesia was quantified every 20 minutes, over a two-hour period, using a scale of 0-4. A score of 0 indicates no abnormal movement, and a score of 4 describes continuous and uninterruptable dyskinetic movements; 12 (4 x 3 body segments) is the maximum score possible for a given time point. Dyskinesia was quantified every other minute during *in vivo* electrophysiology experiments.

### *Ex vivo* Electrophysiology

For excitability, current-response curves (Figure 2F-I) were compared using a one-way repeated measures ANOVA, either across cell-types (Figure 2F) or within a cell-type before and after application of SKF-81297 (Figure 2G-I), with a Tukey. Passive and active properties across the three cell-types were compared using a nonparametric Kruskal-Wallis (KW) test, with a posthoc Tukey test (Table 1) or within cell-types before and after application of SKF-81297 using a paired, nonparametric Wilcoxon signed-rank test (Table 2). Significant p-values were determined following Bonferroni correction for multiple comparisons. Frequency and amplitude of mESPCS (Figure S3A-C), as well as AMPA/NMDA ratio (Figure 4A-C) were compared between the three cell-types using a KW test. Paired-pulse ratio curves (Figure 4C) were compared between the three cell-types using a one-way repeated measures ANOVA, with a posthoc Tukey test. For mEPSC frequency and amplitude measurements, only cells with at least 500 events were included in the. Cumulative probability plots were generated from 500 randomly selected mEPSC events per cell. Changes in excitability in response to acute SKF application were analyzed by comparing a 10-minute baseline period with the value 10-15 minutes after drug application. Average amplitudes of oEPSCs were quantified manually in Igor. A nonparametric Wilcoxon sign-rank (SR) test was used to compare oEPSC amplitude from M1, S1, or thatalmus onto FosTRAP;Ai14;D2-GFP MSNs. In all experiments involving optical stimulation, data was drawn from stimulations at 0.5, 1, 2 ,4 mW, with statistical comparisons being made at 4 mW (Figure 4H,K,N).

***In Vivo* Electrophysiology**

For the majority of analyses of single-unit firing rate and behavior, firing rate was averaged in 1 minute bins. Modulation of firing rate by levodopa was determined by comparing single-unit firing rates before and after drug administration, during the peak behavioral effects. The 30-minute baseline period was compared to a 30-minute period following drug injection (10-40 minutes post-injection). Following levodopa administration, unlabeled single-units were categorized into three broad groups as follows, based on significant changes in firing rate ($p < 0.01$, Wilcoxon rank-sum test (RS)) following levodopa treatment: putative dMSNs (On MSNs, increase in firing rate), putative iMSNs (Off MSNs, decrease in firing rate), or no change units (NC, nonsignificant change in firing rate) (Figure 1G). For levodopa sessions, putative dMSNs were further divided using behavior-based methods, as described previously (Ryan et al, 2018). For the behavior-based method, AIM scores were also averaged in 1 minute bins and correlated with firing rate using linear regression. Labeled TRAPed neurons or putative dMSNs with a significant correlation ($R^2$ > 0.30) to AIM score were labeled dyskinesia (DYSK) units and those with no significant correlation ($R^2$ < 0.35) to AIMs were classified as on-unclassified (ON) units (Figures 1L-M and Supp Figure 1G).

Firing rates of parkinsonian mice before (Park) and after drug administration (levodopa (LID), Figure 1K) were compared between optogenetically labeled TRAPed putative dMSNs and all putative dMSNs using Wilcoxon rank-sum test (RS). Comparisons between the average dyskinesia correlation of optogenetically labeled TRAPed and all putative dMSNs were made using Wilcoxon rank-sum test (RS).

***Monosynaptic Rabies Tracing***

Custom analyses were written in MATLAB for quantification of rabies labeled cells. The relative number of presynaptic neurons was quantified by dividing the total number of presynaptic neurons in the specified brain region by the total number of co-infected (sTpEpB, green and rabies, red) striatal neurons (Figure S2D). The relative proportion of presynaptic neurons was quantified by dividing the total number of presynaptic neurons in the specified brain region by the total number of presynaptic (rabies-labeled, red) extra-striatal neurons detected in the whole brain. Results were when pooled across mice of the same genotype (D1-Cre, A2a-Cre, or TRAP-CreER) and treatment condition (control, levodopa-naïve parkinsonian, or levodopa-treated parkinsonian). Comparisons between treatments conditions for iMSNs and dMSNs were compared using a nonparametric Kruskal-Wallis test, with posthoc Tukey test (Figures 3 and S2). Comparisons between $TRAP^{CreER}$ and D1-Cre levodopa-treated, parkinsonian mice were made using a Wilcoxon rank-sum test (RS).
